# Intra-specific NLR allelic diversity and genomic landscape for plant resistance association studies

**DOI:** 10.1101/2025.08.01.668142

**Authors:** Javier Belinchon-Moreno, Aurélie Berard, Aurélie Canaguier, Isabelle Le-Clainche, Pascale Mistral, Karine Leyre, Vincent Rittener-Ruff, Jacques Lagnel, Damien Hinsinger, Patricia Faivre-Rampant, Nathalie Boissot

## Abstract

The identification of nucleotide-binding domain leucine-rich repeat receptor genes (NLRs) across multiple accessions is essential to capture their genetic diversity within a plant species. Using Nanopore adaptive sampling and *de novo* assembly, we accurately reconstructed the NLR regions across 143 *Cucumis melo* accessions representing diverse botanical groups and geographical origins. NLR annotation evidenced diverse cluster architectures and unexpected variation in NLR content across accessions, leading to unsaturated allelic diversity curves. Using this diversity, we further proposed pan-NLRome graph- and k-mer-based genome-wide association studies (GWAS), which, using Fusarium wilt races 1 and 2 severity data, accurately identified *Fom-1, Fom-2*, and novel non-NLR candidates. Furthermore, we extended these approaches for the identification of a candidate gene for flaccid necrosis caused by zucchini yellow mosaic virus. Our study offers a comprehensive view of NLR diversity in melon, overcoming the limitations of single-reference analyses, and supporting future efforts in NLR-focused GWAS and resistance breeding.

## Introduction

Plants rely on a dual-branch innate immune system to combat infections that co-evolves with pathogens^1^. The initial branch uses transmembrane pattern recognition receptors (PRRs) to detect conserved molecules shared by various microbial classes, as well as intrinsic molecules that result from cellular injury^1,2^. To overcome the first layer of defense, pathogens often deliver effector molecules into plant cells, which are recognized by intracellular receptors, triggering a second layer of resistance that is frequently stronger^1,3^. Most of these intracellular receptors are nucleotide-binding domain (NBD) leucine-rich repeat (LRR) receptor genes (NLRs)^4^. Despite their highly polymorphic nature^5^, NLRs typically present a conserved intrinsic structure comprising three domains with distinct and variable functions^3,6^. The N-terminal domain is usually a Toll/interleukin-1 receptor/resistance protein (TIR), a coiled-coil (CC), or a resistance to powdery mildew 8 (RPW8)-like domain^3^. Additionally, truncated NLRs missing partial or complete domains can remain functional^4,7,8^, broadening the range of possible configurations and immune strategies.

Given the extreme intraspecific diversity and frequent presence-absence (P/A) polymorphic variation of NLR regions^9–12^, reference genomes likely contain a small part of the species NLR diversity, making impossible the use of resequencing for its complete and accurate characterization^4,10^. The assembly of NLR regions from multiple genotypes stands as a great solution, but the frequent organization of NLRs into complex clusters with high copy-number variation, often enriched with repetitive elements^4,13,14^, makes these regions challenging to assemble. Despite this, pan-genomes for several wild and cultivated plant species have been released^9,12,15–18^, enabling comprehensive analyses of their pan-NLRomes. Furthermore, some targeted pan-NLRomes have also been published^10,11,19–22^, mainly using the resistance gene enrichment sequencing method (RenSeq) combined with long read technology. However, most of these studies focused on evolution, lacking investigation into the links between specific NLR alleles and resistance phenotypes. Genome-wide association studies (GWAS) using reference-free approaches based on pan-genomes have been proposed^23–25^, potentially enabling the identification of genomic variation patterns associated with important phenotypic traits. These methods included the construction of variation graphs, which enable accurate genotyping of variants ranging from single-nucleotide polymorphisms (SNPs) to structural variants (SVs)^26–29^, as well as the use of defined short sequences (k-mers) that serve as markers for the P/A of variants of all sizes^30–32^. Nevertheless, the application of these approaches remains relatively limited in practice.

Here, we employed Nanopore adaptive sampling (NAS) to target the NLR regions of 143 melon accessions. We annotated NLRs across assemblies and conducted their classification, characterization and diversity analysis. Finally, we proposed two novel approaches that move beyond traditional single-reference GWAS: one leveraging a graph-based pan-NLRome as reference, and the other using a fully reference-free k-mer method.

## Results

### NANOPORE ADAPTIVE SAMPLING ENABLED HIGH-QUALITY TARGETED ASSEMBLY OF MELON NLR REGIONS

We sequenced with NAS and successfully assembled 21 genomic regions likely containing NLRs (Fig. 1a) across 143 melon accessions, representing diverse geographical origins and botanical groups (Supplementary Table 1, Extended Data Fig. 1). A detailed summary of sequencing metrics, reads filtering and assembly steps is provided in Supplementary Tables 2 and 3. The average sequencing depth across all *de novo* assembled contigs was of 41.97×. Ninety-five percent of contigs presented a sequence depth over 14×, and over 63% had a sequence depth greater than 30× (Supplementary Table 4). We estimated accuracy and completeness of the assembled regions by quantifying mismatches observed when mapping the reads used for assembly, as in ref. ^10^. The average quality across assembled contigs for all the accessions exceeded Q30. Ninety-five percent of the contigs had a quality above Q22, with more than 51% surpassing Q30 (Supplementary Table 4). Regarding assembly contiguity, the vast majority of NLR regions were assembled into a single contig. Only one region was fragmented into three contigs, while just 2.71% of the regions were split into two contigs (Supplementary Table 5).

**Figure 1.**
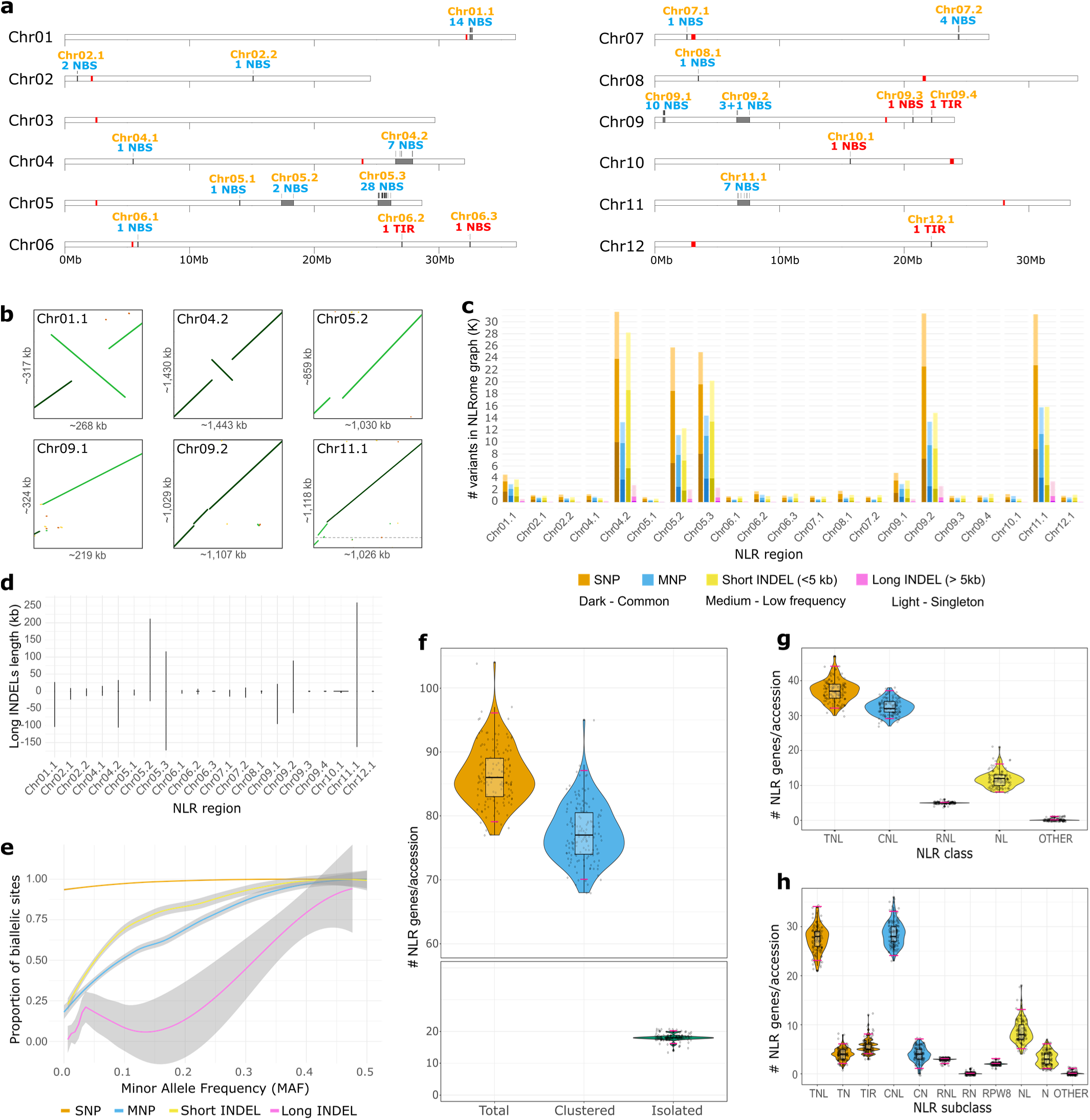
Nucleotide- and NLR-based observed diversity of the pan-NLRome. **a,** Predicted NLRs distribution in the ANSO-77 reference genome assembly. Red vertical bars represent predicted centromeres using the quarTeT software v. 1.1.6^37^ with default settings and adding the repeat annotation and gene annotation files. Gray vertical bars represent NLR regions, labelled in blue when used for adaptive sampling, and labelled in red when reference-guided assembled. **b,** Structural variations detected through the assembly of multiple accessions compared to ANSO-77 (x-axis). **c,** Number of variants in each pan-NLRome subgraph, classified by variant type (color) and frequency (lightness) across the studied accessions. **d,** Size distribution of INDELS >50bp per pan-NLRome subgraph. Data is represented in the shape of violin plots, with most of the size observations being unique. **e,** Pan-NLRome MAF spectrum of the proportion of biallelic sites for each variant type (SNP, MNP, short, and long INDEL). Smoothed lines were fitted using the locally estimated scatterplot smoothing (LOESS) method with the ggplot2 R package^38^. Gray shaded areas around the fitted lines represent their 95% confidence interval. For c, d and e, variants were extracted from the pan-NLRome graph using ANSO-77 as the reference path. Each subgraph was constructed with 143 to 162 pseudo-haplotypes (see Materials and Methods). **f,** Total, clustered and isolated NLR content distribution across the 143 accessions. **g,** NLR content distribution per general NLR class across the 143 accessions. **h,** NLR content distribution per specific NLR subclass across the 143 accessions. In f, g and h, the path with the maximum number of NLRs possible was considered when more than one pseudo-haplotype was assembled for a single NLR region. Solid black horizontal lines represent medians; solid pink horizontal lines represent Bayesian 95% highest density intervals (HDIs); small black circles around the box-plots represent individual observations.

### THE ASSEMBLY OF NLR REGIONS REVEALED HIGH STRUCTURAL DIVERSITY

We generated dot plots of the NLR regions of 143 accessions against the regions of ANSO-77. We observed a great diversity of SVs, some of them shared across multiple accessions (Fig. 1b, Supplementary Table 3). Particularly, we visually identified two inversions of ∼150 and ∼250 kb, two ∼80-130 kb insertions/deletions (INDELs), and a large ∼860 kb INDEL (Fig. 1b). This long INDEL, which was present in 37 accessions but absent in ANSO-77, could not have been fully captured using NAS due to its long size^33^, reducing assembly contiguity of the zone.

We constructed a pan-NLRome variation graph comprising 21 sub-graphs to comprehensively characterize and explore the diversity captured in the assembled NLR regions. Each sub-graph consisted of nodes and edges, where each node represented a DNA segment, and haplotype sequences were depicted as walks or paths through the graph^34^. The pan-NLRome graph presented a total size of 154.77 Mb, with 949,683 nodes and 1,314,842 edges (Supplementary Table 6). The number of nodes and edges across the 21 sub-graphs was closely linked to the region size (Extended Data Fig. 2a). The sub-graphs showed a compression factor (sequence length included in the graph divided by the graph size) ranging from 9.4 (for region Chr10.1) to 103.8 (for region Chr06.1), with an average of 68.5 (Extended Data Fig. 2b).

We extracted variants from all the 143 assemblies compared to a reference, ANSO-77. We identified 241,354 variant sites and 384,442 alternative alleles (Supplementary Table 6). Around 46% of the variant sites were top-level snarls, with ∼41% nested inside top-level snarls (Supplementary Table 6), which was expected given the complexity of NLR regions. We evidenced the particular complexity of region Chr05.3, as it presented 93 very complex snarls of level five and four of level six (Supplementary Table 6). Globally, 18% of the variant sites were multiallelic (Extended Data Fig. 2c, Supplementary Table 6). Most variants were SNPs and short INDELs present in less than 10% of accessions, with a significant proportion classified as singletons (Fig. 1c). We observed in general longer INDELs inside larger NLR regions, most of them being unique (Fig. 1d).

The proportion of biallelic sites is a key factor for GWAS, as filtering multiallelic variants and applying a MAF threshold of 0.05 is a common practice^35,36^. We assessed the impact of minimum allele frequency (MAF) for each variant type on the proportion of biallelic sites (Fig. 1e). SNPs remained predominantly biallelic regardless of the selected MAF, but applying a MAF threshold of 0.05 would result in the loss of approximately 80% of long INDELs, and around 50% of both MNPs and short INDELs (Fig. 1e).

#### TNLs AND CNLs ARE PREDOMINANT WITHIN THE REDUCED REPERTOIRE OF NLR GENES IN MELON

Detailed gene annotation metrics for all assembled accessions are summarized in Supplementary Table 7. We identified 13,330 NLRs across the 143 accessions using four complementary tools. Since some regions in certain accessions were assembled into two pseudo-haplotypes, we estimated both the minimum and maximum possible NLR counts on these accessions based on their pseudo-haplotype structures (Extended Data Fig. 3). We considered the maximum possible NLR count per accession for consistency across the study, which ranged from 77 to 104 (Fig. 1f). We defined NLR clusters as gene groups separated by less than 1.5 Mb. Across accessions, 87–91% of NLRs were located within the defined clusters. Each accession contained 7–10 isolated NLRs, while the number of clustered NLRs ranged from 68 to 95 (Fig. 1f).

The most frequent NLR class was TNL, closely followed by CNL (Fig. 1g), due to the inclusion of isolated TIR domains (Fig. 1h). In fact, the content of canonical CC-NBS-LRR genes was slightly higher than that of TIR-NBS-LRR genes (Fig. 1h). The third class based on NLR count was NLs, while RNLs were less abundant and showed little variability in their numbers across accessions (Fig. 1g).

### NLR CLASSES ARE NOT MIXED WITHIN THE MELON GENOME

We identified NLRs across the 143 accessions in 20 out of the 21 assembled NLR regions, with no NLRs found in region Chr04.1, excluded for subsequent analyses. Five small-sized NLR regions, typically containing isolated NLRs, contained no NLRs in the assembly of some accessions (Fig. 2). Particularly, we only annotated NLRs in Chr10.1 in four accessions. No accession lacked NLRs in more than two of the 20 identified NLR-containing regions. We observed NLRs on all chromosomes except on Chr03 (Fig. 1a). The distribution of NLRs was uneven among chromosomes, with Chr05 accounting for approximately 30% of them (Fig. 2). We also observed a higher number of NRLs on Chr01, Chr09, Chr11, and Chr04 compared to the other chromosomes.

**Figure 2.**
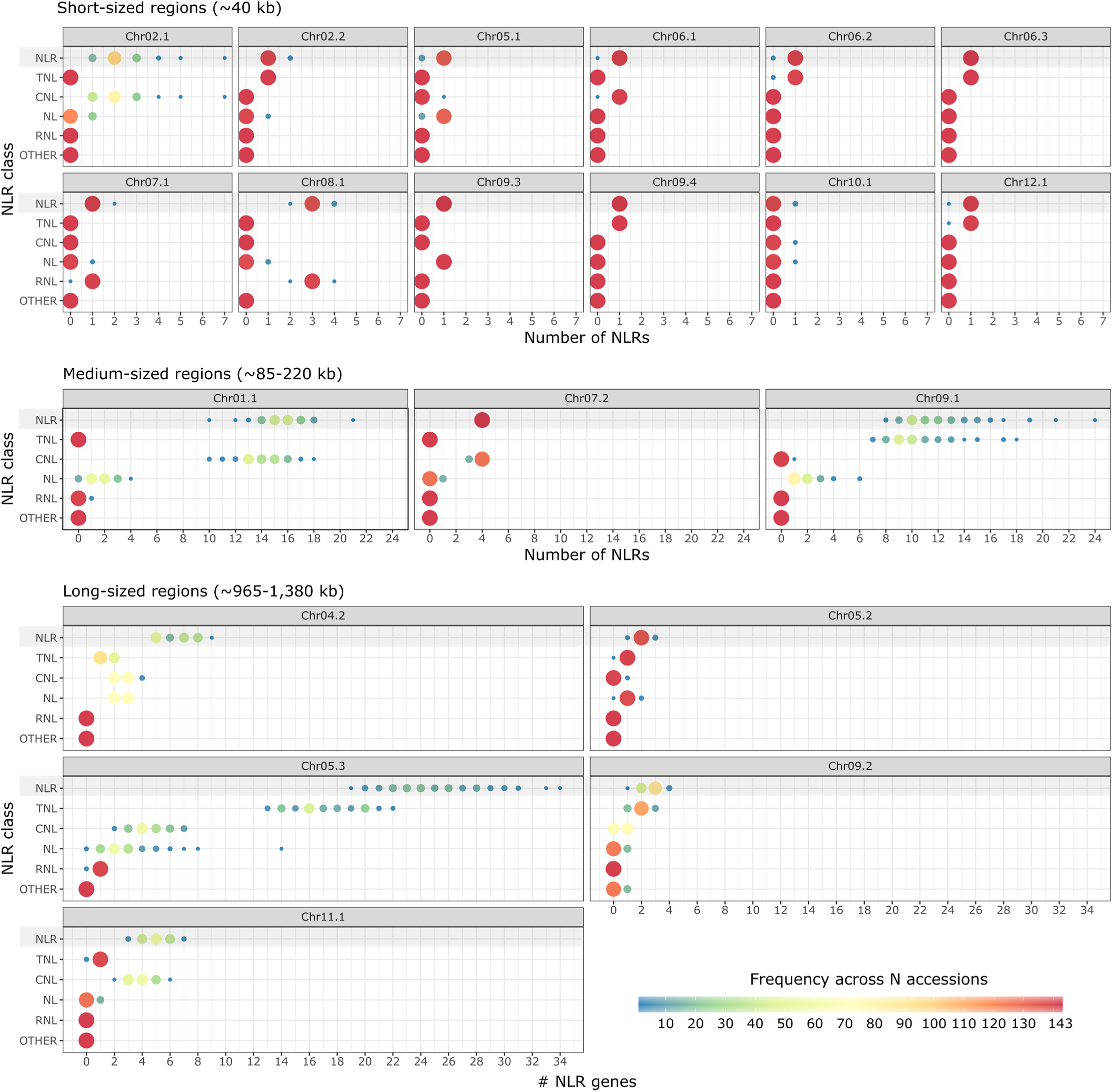
Patterns of NLR copy-number variation among NLR regions and NLR classes. The color and size of the bubbles represent the number of accessions containing a given NLR count. NLR regions are categorized into three size classes: short (≈40 kb) at the top, medium-sized (≈85–220 kb) in the middle, and long-sized (≈965–1,380 kb) at the bottom. When more than one pseudo-haplotype was assembled for a single NLR region and accession, the path with the maximum number of NLRs possible was considered.

Overall, NLR regions differed drastically from each other in terms of length, span and average NLR content. We divided the 20 NLR regions in 12 short- (∼40 kb), three medium- (∼85-220 kb) and five long-sized (∼965-1,380 kb) (Fig. 2). Short-sized regions primarily harbored isolated NLRs, except regions Chr02.1 and Chr08.1, for which 68.5% and 95.8% of accessions contained two and three NLRs, respectively. Medium-sized regions displayed variable NLR content. Region Chr07.2 contained four NLRs in all accessions, while regions Chr01.1 and Chr09.1 harbored at least 10 and eight NLRs, respectively. The same trend was observed in long-sized regions. Notably, cluster Chr05.2 averaged only two NLRs, while region Chr05.3 included at least 19. In terms of copy number variability, we observed four highly variable NLR regions across accessions: Chr02.1 (range 1-7), Chr01.1 (10-21), Chr09.1 (8-24), Chr04.2 (5-9), Ch05.3 (19-34) and Chr11.1 (3-7) (Fig. 2)

Lastly, the classification of NLRs within the defined genomic regions revealed distinct patterns of NLR class composition. In short- and medium-sized regions, the majority of detected NLRs belonged to a single class (Fig. 2). Among those harboring clustered NLRs, we observed three CNL-dominated clusters (Chr02.1, Chr01.1, and Chr07.2), a distinct RNL-enriched cluster (Chr08.1), and a high-density TNL cluster (Chr09.1) (Fig. 2). Long-sized regions commonly exhibited successions of NLR blocks belonging to the same class. For instance, examination of annotations across all analyzed accessions for cluster Chr11.1 revealed the pattern CNL-TNL- CNL, separated by ∼25 and ∼250 kb, respectively. Similarly, Chr05.3 exhibited a pattern RNL- CNL-TNL, separated by ∼300 and ∼40 kb, respectively.

### THE MELON PAN-NLROME NEEDS MORE THAN 143 GENOMES OF WORLWIDE-DISTRIBUTED ACCESSIONS TO REACH SATURATION

We first evaluated the growth and saturation dynamics of the pan-NLRome sub-graphs. In some sub-graphs, such as those of regions Chr09.1, Chr09.2, Chr10.1, and Chr11.1, more than half of the sequences were shared by only one or two haplotypes (Fig. 3a). In contrast, in those of regions like Chr04.2, Chr05.1, Chr06.1, Chr06.3, and Chr08.1, more than half of the sequences were shared by more than half of the haplotypes (Fig. 3a).

**Figure 3.**
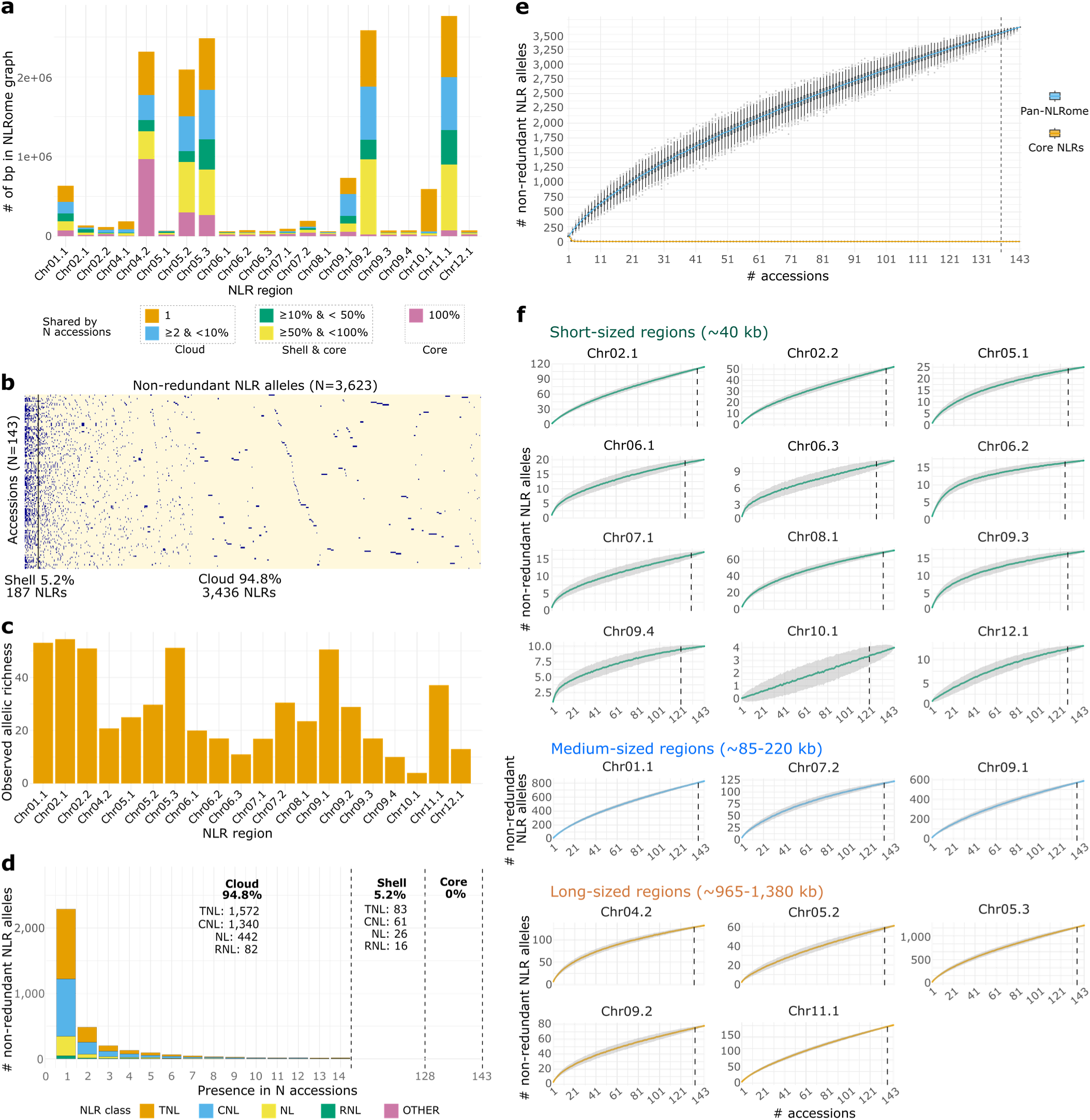
Nucleotide- and allele-based saturation analysis of the pan-NLRome. **a,** Final step of the growth saturation curves showing the cumulative sequence size across the 21 pan-NLRome subgraphs. Saturation growth was obtained from Panacus^42^. Each subgraph was constructed with 143 to 162 pseudo-haplotypes (see Materials and Methods). **b,** Heatmap representing the variation patterns of nr NLR alleles across the 143 accessions. Blue squares represent nr NLR alleles present in the corresponding accession, while yellow squares represent those absent. A black vertical line divides nr alleles belonging to the cloud (<10% of accessions) and shell (10%≤N<90%) pan-genome. **c,** Observed allelic richness per NLR region. The allelic richness in a given NLR region was calculated as the number of nr alleles accounted in that region divided by its average number of NLR loci across the 143 accessions. **d,** Distribution of the number of nr alleles shared by N accessions among different NLR classes in the cloud pan-genome. The percentage of nr alleles belonging to the cloud, shell and core pan-genome is shown at the top, with the detailed number of alelles per class displayed below it. **e,** Saturation growth curve of the variation in the number of nr alleles within the pan-NLRome and core NLRome, considering all NLR regions. Boxplots were fitted at each given number of accessions, with a colored line connecting medians. Gray dots mark outliers beyond 1.5 x Interquartile Ratio, and whiskers cover all non-outlier values. **f,** Saturation growth curve of the pan-NLRome showing changes in the number of nr alleles across individual NLR regions. Colored lines connect means for each given number of accessions. The shaded areas around the lines mark one standard deviation around the means. For e and f, saturation curves were generated by incrementally increasing the number of accessions (from 1 to 143), selecting nr alleles present at each step, and performing 1,000 permutations per data point. Black, dotted, vertical lines mark the minimum number of accessions needed to obtain 95% of the observed nr alleles.

Annotated NLRs were classified into 69 orthogroups (Supplementary Table 8). Pan-NLRome saturation analysis based on orthogroups showed 95% saturation with 96 randomly chosen accessions (Extended Data Fig. 4). Because NLR allelic variants or paralogs can trigger distinct immune responses against different pathogens^39–41^, and because both susceptible and resistant alleles of the same NLR typically reside within the same orthogroup, orthgroup-based clustering is not informative for dissecting disease resistance. Therefore, we established the P/A patterns of 3,623 non-redundant (nr) NLR alleles across all the accessions (Fig. 3b). In general, allelic richness of NLR regions (Fig. 3c) was related to NLR count (Fig. 2) except for region Chr02.2, mostly containing a single isolated TNL across most accessions (Fig. 2) but displaying one of the highest allelic richness values (Fig. 3c). We assigned nr NLR alleles to three groups: cloud (allele present in less than 10% of the genomes), shell (allele present in 10-90% of the genomes), and core pan-genome (allele present in more than 90% of the genomes). Ninety-five percent of nr alleles were part of the cloud group, of which 63% were singletons (Fig. 3d), while no allele was found in the core group. Regarding NLR classes, TNLs, CNLs and NLs were distributed among cloud and shell following the general trend (Fig. 3d). However, RNLs were more frequent than expected in the shell pan-genome (16.3%) (Fig. 3d). The most conserved allele, present in 120 accessions, was an isolated TNL in region Chr12.1. Global pan-NLRome growth curve did not reach a plateau (Fig. 3e), indicating that the discovery of NLR alleles remained far from saturation. We observed the same trend for each independent NLR region, whatever its size and location (Fig. 3f).

### A FOCUS ON THE EXPERT-ANNOTATED CNL BLOCK OF CHR05.3 CONFIRMED THE LACK OF NLR DIVERSITY SATURATION

A CNL cluster spanning the *Vat* genes and included inside the region Chr05.3 has been well characterized^43,44^. We were able to manually annotate *Vat* alleles here, providing significantly higher accuracy and confidence than the automated annotation approaches used for the rest of NLR regions. The region sized between 240 and 495 kb in the accessions panel. We globally annotated 774 *Vat* alleles (167 nr), with a number of *Vat* across accessions ranging from two to nine (considering the haplotype with the maximum number of *Vat* alleles when two were available) (Fig. 4a, Extended Data Fig. 5). The growth and saturation dynamics of this expert- annotated region confirmed the lack of saturation of nr alleles in the panel (Fig. 4b).

**Figure 4.**
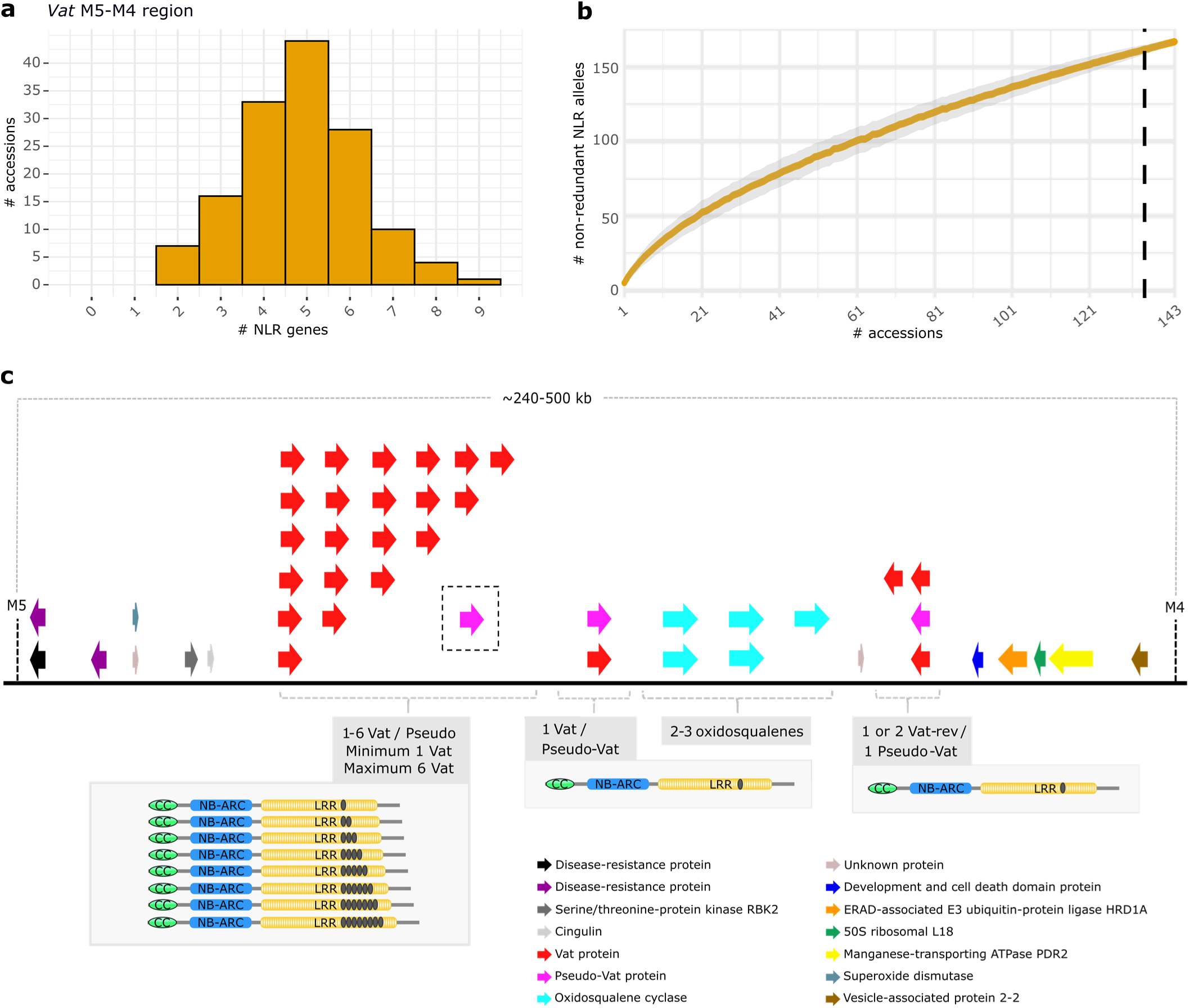
Observed diversity of the Vat NLR cluster in 143 accessions. **a,** Patterns of NLR copy-number variation in the Vat region. When more than one pseudo-haplotype of Vat was assembled for an accession, the path with the maximum number of NLRs possible was considered. **b**, Saturation growth curve showing changes in the number of nr NLR alleles in the Vat region. The curve was generated by incrementally increasing the number of accessions (from 1 to 143), selecting nr Vat alleles present at each step, and performing 1,000 permutations per data point. The orange line connects means for each given number of accessions. The shaded area around the line mark one standard deviation around the means. Black, dotted, vertical lines mark the minimum number of accessions needed to obtain 95% of the observed nr alleles. **c**, Canonical structure of the Vat region. The consensus for Vat gene content was established based on 156 pseudo-haplotypes across 143 accessions. The consensus for the remaining genes within the Vat M5-M4 region was derived from the accessions annotated in ref. ^44^. Black ellipses within the NLRs indicate the number of R65aa motifs present in the LRR2 domain. The pseudo-Vat gene enclosed by a dashed square denotes the potential presence of one or more pseudo-Vat genes among the 2 to 8 Vat homologs identified in this region.

Aside copy-number variation, the number of leucine-rich repeat sequences of 65 amino acids (R65aa) that *Vat* alleles contain in the LRR2 part is a key factor of *Vat* resistance specificity^43^. Most observed *Vat* alleles presented one to eight R65aa motifs, with extreme cases presenting 16 and 18 R65aa motifs (Supplementary Table 9). One to five and seven R65aa were very common, with more than 60 *Vat* alleles annotated in each category (Supplementary Table 9).

We consistently observed one to six functional *Vat* homologs across accessions and one or two *Vat* reverse-transcribed (*VatRev*). We analyzed 156 haplotypes to depict the “canonical” structure of the *Vat* cluster (Fig. 4c, Supplementary Table 10), in which we observed 126 combinations of functional *Vat* homologs defined by their number of R65aa motifs (Supplementary Table 10) and 135 combinations of nr *Vat* alleles (Supplementary Table 10).

### PAN-NLROME DIVERSITY PREDICTION SUGGESTED BOTANICAL GROUPS AND ORIGINS WITH HIGH POTENTIAL FOR INMUNITY EXPLORATION

As our results showed that any of the 20 NLR regions reached saturation after the inclusion of 143 accessions, we investigated alpha diversity estimators to correct for alleles that might be present but not detected in our sample due to limited sample size. Using an analogy to the classical ecological definition of alpha diversity, nr NLR alleles were equated to species within a functional community, and NLR regions to a defined ecological unit, such as a community. The number of nr alleles in each NLR region of each accession is therefore analogous to the number of species in a given ecological sample. We calculated three indices of alpha diversity that account for richness and evenness at different levels: the Chao index, the Shannon index and the Simpson index. Chao2 estimates indicated that between 3,960 and 8,521 nr NLR alleles were still undiscovered, with five regions potentially missing two thirds of their complete diversity (Fig. 5a). Observed and estimated Shannon and Simpson indices, both giving less weight to less abundant NLR alleles than Chao, exhibited less deviation than this last (Fig. 5b). This pattern was expected, as we previously observed a high number of singletons in our data (Fig. 3d). Then, we constructed sample completeness curves (Fig. 5c) describing how the estimated sample completeness (i.e., the proportion of total diversity captured) changes as a function of sample size. In some regions, such as Chr06.2, Chr09.3 or Chr09.4, the observed accessions were sufficient to approach nearly 100% sample completeness while for others, particularly those with high Chao2 estimate such as Chr01.1, Chr05.3 or Chr09.1, coverage remained near 75%, suggesting again a large number of undetected nr NLR alleles. Extrapolation of these curves indicated that more than twice of the current number of accessions would be required to surpass 90% completeness (Fig. 5c).

**Figure 5.**
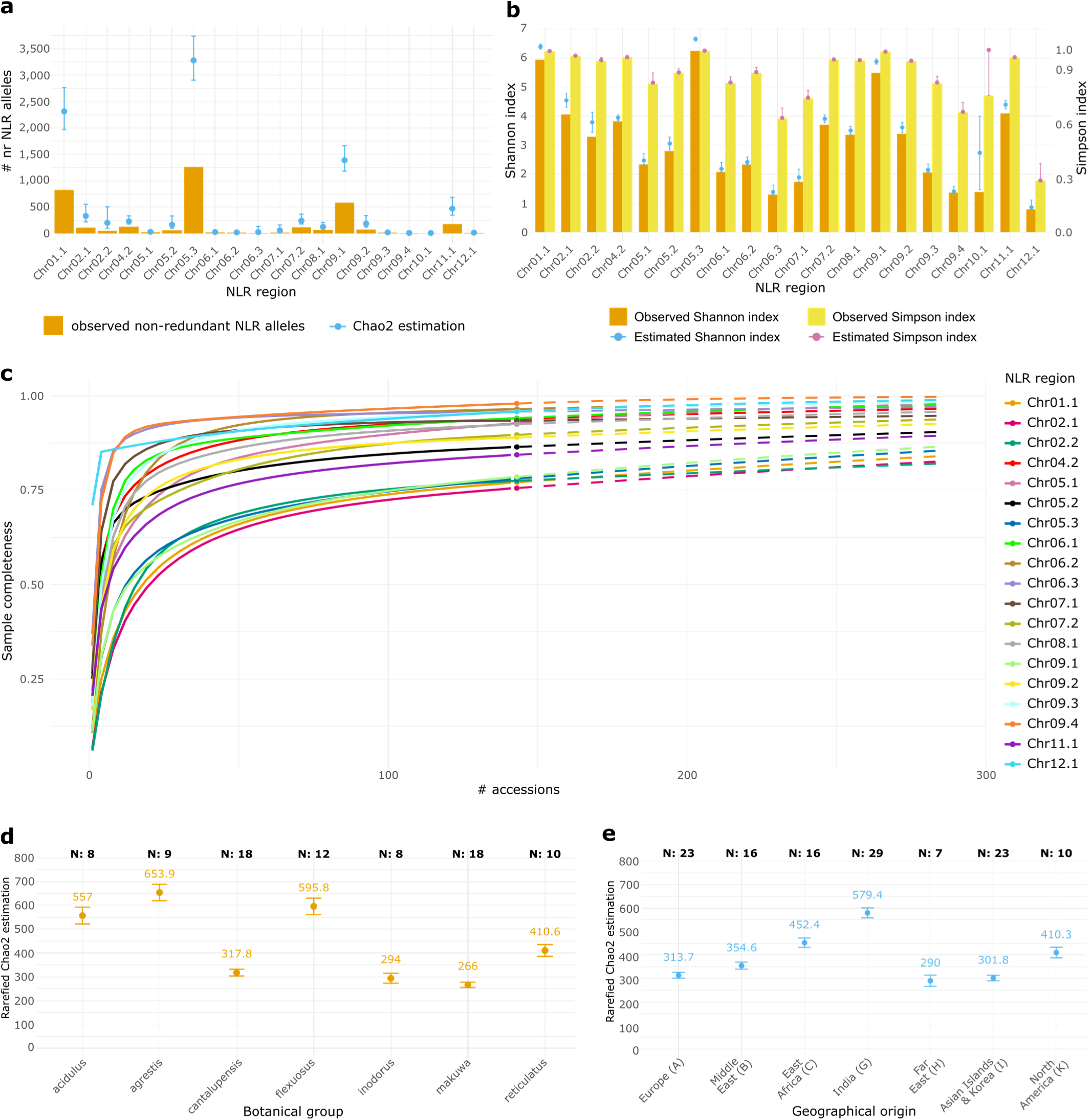
Estimated diversity of the pan-NLRome in melon. All estimations were performed with the R package iNEXT^46^**. a,** Observed nr alleles in 143 accessions and Chao2 estimates in the whole species per NLR region. Orange bars represent the number of observed nr alleles across NLR regions. Blue points represent the asymptotic Chao2 index estimates per NLR region. Blue vertical lines represent the 95% confidence intervals for the Chao2 estimates. **b,** Observed and estimated Shannon and Simpson diversity indices across NLR regions. Bars represent the observed indices per NLR region within 143 accessions and points represent index estimates in the melon species. Blue and pink vertical lines represent the 95% confidence intervals for the Shannon and Simpson estimates. **c**, Estimation of number of accessions needed to saturate the melon pan-NLRome. Solid lines represent observed data, while dashed lines depict estimates from these observations. Region Chr10.1 was excluded as not enough data was available. **d and e,** Chao2 diversity estimates for botanical (d) and geographical (e) subgroups. The sample size per subgroup is included at the top of the graphs. As the sample size among subgroups was different (eight to 18 for botanical groups, and 7 to 29 for geographical origins), Chao2 estimates were calculated by rarefaction to the smallest sample size. Only subgroups with more than seven observations were included in the analysis.

Finally, we explored the botanical and geographical components of the diversity predictions by calculating rarefied Chao2 estimates, standardized to the smallest sample size among groups^45^. *Acidulus* and *flexuosus* groups exhibited high rarefied Chao2 index, aligning with their higher NLR content (Fig. 5d, Extended Data Fig. 6). Accessions belonging to these groups mostly come from India (Fig. 5e; Supplementary Table 1). Notably, *agrestis* accessions presented the highest rarefied Chao2 index (Fig. 5d) while harboring a lower median number of genes than *acidulus* and *flexuosus (*Extended Data Fig. 6), highlighting their highest allelic diversity.

### GRAPH- AND K-MER-BASED GWAS PROVIDED IMPROVED ACCURACY AND REVEALED ADDITIONAL CANDIDATES BEYOND THE SNP-BASED METHOD

Most commonly used GWAS rely on mapping short reads to a single reference genome, followed by genetic association analyses based on SNPs or small INDELs^47^. We compared whole-genome SNP-based GWAS to graph-based and k-mer-based GWAS, using the resources generated in this study. We benchmarked both methods using symptoms severity data to Fusarium wilt races 1 and 2, mainly controlled by two dominant major cloned genes *Fom-2* (resistance to races 0 and 1)^48^ and *Fom-1* (resistance to races 0 and 2)^40^.

Broad sense heritability values, calculated for 126 and 127 accessions, were of 94.9 and 91.1% for Fusarium wilt race 1 and 2, respectively, validating the experimental protocols. We extracted Best Linear Unbiased Predictors (BLUPs) from the scoring phenotypic data (Supplementary Table 11). Using the STRUCTURE algorithm on the SNP matrix complemented with the Evanno group selection method, we inferred five groups as the most relevant genetic organization (Extended Data Fig. 7a,b; Supplementary Table 11). For whole-genome SNP-based GWAS, we selected the model incorporating both kinship and five population structure groups for both phenotypes, as it provided the best fit based on Q-Q plots inspection (Extended Data Fig. 7c-e). We selected the model including only the kinship for both graph- and k-mer-based approaches, as NLR-targeted structure estimation could be biased due to the high number of copy-number variations (CNVs) highlighted in previous paragraphs.

For Fusarium race 1, as expected, we obtained significant SNP peaks over the established Bonferroni thresholds around the NLR cluster containing the cloned *Fom-2* (cluster Chr11.1) (Extended Data Fig. 8a). Linkage disequilibrium (LD) analysis around the top-linked SNP suggested a quantitative trait locus (QTL) of 3.88 Mb that included 20 other significant SNPs, with the lowest bound located ∼2.2 Mb downstream from the *Fom-2* homolog (Fig. 6a). We observed a second QTL of 146.88 kb including five other significant SNPs and located ∼3.5 kb downstream from *Fom-2*. Graph-based GWAS yielded 85 nodes above the defined significance threshold. Some of these nodes corresponded to variants of region Chr11.1, with a second group of associated signals in Chr05.3 region (Extended Data Fig. 8a), not detected in SNP-based GWAS. K-mer-based GWAS on targeted NLR regions yielded 577 k-mers over the defined Bonferroni significance threshold. Local assembly of groups of these k-mers sharing identical p-values provided 30 assembled sequences that mapped on the ANSO-77 whole-genome assembly forming a pronounced peak in the Chr11.1 region and, interestingly, another peak overlapping the region Chr05.3. Focusing on the *Fom-2* regions of ANSO-77 (susceptible) and Isabelle (resistant) as example, we observed that many of the top-associated nodes and k-mers located within or in very close proximity to *Fom-2.* Particularly, the top k-mer block belonged to the sequence of Isabelle’s *Fom-2* (Fig. 6a,b). Further investigation on the signals on Chr05.3 region revealed two gene candidates that were not annotated as NLRs by our costumed pipeline (Fig. 6e,f). Protein BLAST against the nr protein sequences database suggested that these genes belong to the GDSL family of serine esterases/lipases, already associated in many studies with defense responses, including Fusarium head blight resistance in wheat^49^ or Fusarium wilt resistance in bottle gourd (*Lagenaria siceraria*)^50^.

**Figure 6.**
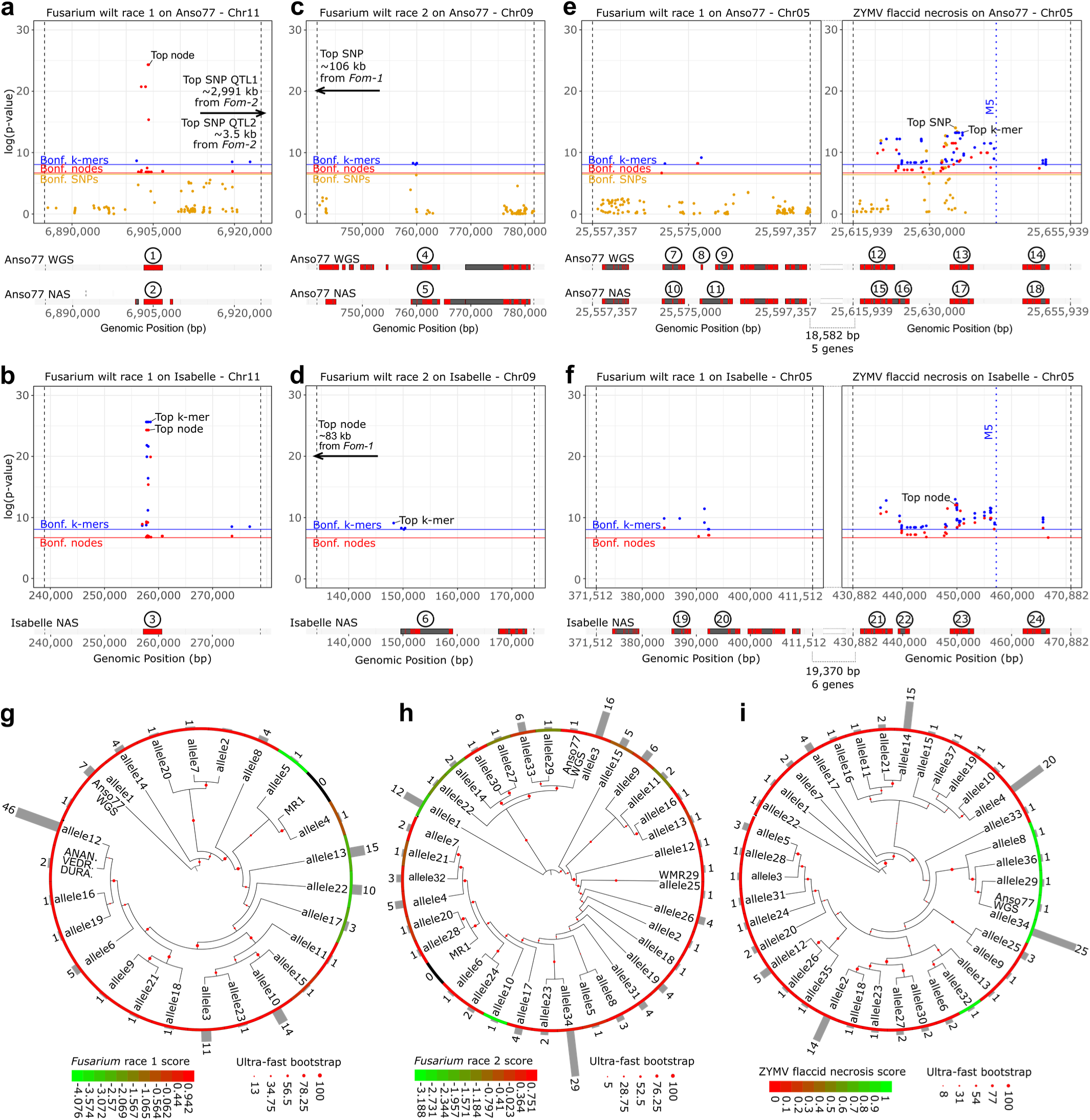
SNP-, graph- and k-mer-based GWAS results for Fusarium wilt races 1 and 2, and ZYMV flaccid necrosis, and post-GWAS analyses. **a,b,** Focus on Fom-2. c,d, Focus on Fom-1. **e,f**, Focus on region Chr05.3 for Fusarium race 1 (left) and ZYMV flaccid necrosis (right). **a,c,e,** GWAS on ANSO-77 (susceptible to Fusarium races 1 and 2, and showing flaccid necrosis). Annotated genes are plotted under the Manhattan plots for the WGS assembly (top) and NAS assembly (bottom). Red blocks represent exons, and black blocks represent introns. **b,d,f,** GWAS on Isabelle (resistant to Fusarium races 1 and 2, and not showing flaccid necrosis). Annotated genes in the NAS assembly are plotted under the Manhattan plots. Red blocks represent exons, and black blocks represent introns. SNPs, graph nodes and k-mer Bonferroni thresholds (α=0.05 corrected by the estimated number of independent variants) are plotted with colored horizontal lines. Genes ID are as follows: 1, _Chr11_001675 (reannotated Fom-2); 2, ME00509#1#utg131_Chr11.1_000038 (reannotated Fom-2); 3, ME00791#1#utg22_Chr11.1_000033 (Fom-2); 4, _Chr09_001533 (reannotated Fom-1); 5, ME00509#1#utg284_Chr09.1_000017 (reannotated Fom-1); 6, ME00791#1#utg66_Chr09.1_000017 (reannotated Fom-1); 7, Chr05.927; 8, Chr05.928; 9, Chr05.929; 10, ME00509#1#utg45_Chr05.3_000092; 11, ME00509#1#utg45_Chr05.3_000093; 12, _Chr05.000585 (Candidate RUN1-like); 13, _Chr05.000943 (Candidate RPV1-like); 14, _Chr05.000942; 15, ME00509#1#utg45_Chr05.3_000097 (Candidate RUN1-like); 16, ME00509#1#utg45_Chr05.3_000098; 17, ME00509#1#utg45_Chr05.3_000021 (Candidate RPV1- like); 18, ME00509#1#utg45_Chr05.3_000020; 19, ME00791#1#ctg67_Chr05.3_000031; 20, ME00791#1#ctg67_Chr05.3_000032; 21, ME00791#1#ctg67_Chr05.3_000036 (Candidate RUN1-like); 22, ME00791#1#ctg67_Chr05.3_000037; 23, ME00791#1#ctg67_Chr05.3_000086 (Candidate RPV1-like); 24, ME00791#1#ctg67_Chr05.3_000085. g,h,i, Phylogenetic trees of Fom-2 (g), Fom-1 (h) and RPV1-like candidate (i) alleles identified through 126, 127 and 121 accessions. We also included Fom-2 and Fom-1 resistant and susceptible publicly available alleles from refs. ^40,48^ and those from ANSO-77 WGS. Trees were constructed using maximum likelihood inference with 10,000 ultra-fast bootstraps and 10,000 SH-like approximate likelihood ratio tests. Red to green colors represent the average BLUPs of accessions carrying each allele. Gray bars give the number of accessions carrying each allele.

Regarding Fusarium wilt race 2 data, SNP-based GWAS revealed one of the significant peaks in the Chr09.1 region (Extended Data Fig. 8b). Its top SNP was located ∼106 kb upstream of the *Fom-1* homolog of ANSO-77 (Fig. 6c) but its QTL overlapped the gene. Graph-based GWAS provided two significant nodes that located ∼83 kb upstream from *Fom-1* (Fig. 6d). K- mer-based GWAS yielded 44 signficant k-mers assembled into eight blocks. The top four mapped either 1,265 bp upstream from the *Fom-1* or tagged directly the gene (Fig. 6d).

These results demonstrated that both alternative GWAS were more accurate than the SNP-based method to detect disease-linked NLRs. Moreover, they allowed to detect non-NLR genes located within the NLR regions. The availability of NAS assemblies allowed to examine the diversity of *Fom-2* and *Fom-1* alleles across the 126 and 127 accessions included in GWAS. Notably, Helixer misannotated *Fom-2* in 13 accessions, including the WGS of ANSO-77. To correct this, we re-annotated the gene with the *Fom-2* allele from the resistant accession Isabelle as reference. We obtained four resistance-associated alleles, two of them present in more than 85% of the resistant accessions, and 19 susceptible alleles, with allele12 overrepresented (Fig. 6g). For Fusarium race 2, Helixer systematically misannotated the gene across all the accessions. We therefore performed manual re-annotation using the published *Fom-1* sequence from the resistant cultivar MR-1^40^ as reference. Among the detected alleles, we classified seven as resistance-associated homologs, while 27 conferred susceptibility (Fig. 6h). Notably, allele1 emerged as the most prevalent resistance-associated variant in our panel (Fig. 6h).

### SNP-, GRAPH-, AND K-MER-BASED GWAS CONVERGED ON TWO CANDIDATE LOCI FOR NOVEL ZYMV RESISTANCE

We extended the comparison of SNP-, graph- and k-mer-based GWAS to phenotyping data on plant response to Zucchini Yellow Mosaic Virus (ZYMV) strain E3, which induces flaccid necrosis in some accessions. Flaccid necrosis is a rapid wilting often leading to plant death, potentially associated to extreme hypersensitive response (HR). We employed binomial phenotypic data on a set of 121 accessions for GWAS (Supplementary Table 11). SNP-based GWAS provided a single and very well supported significant peak in chromosome 5, coinciding with the Chr05.3 region (Extended Data Fig. 8c). Graph-based GWAS yielded 68 top- associated nodes, all of them representing variants from the Chr05.3 region (Extended Data Fig. 8c). K-mer-based GWAS produced 1,409 k-mers over the Bonferroni significance threshold, assembled into 112 sequence blocks, all tagging region Chr05.3. We examined the top- associated variants in detail, using ANSO-77 and Isabelle as representative accessions for flaccid necrosis or not (Fig. 6e,f). Top-associated SNP, graph node and k-mer overlapped the same gene, annotated as a TNL. BLAST analysis of this gene against the nr protein database classified it as an RPV1-like protein. We also identified the three types of variants overlapping a gene RUN1-like (Fig. 6e,f). Protein alignment of the RPV1-like candidate across the 121 accessions evidenced 37 different alleles, with four of them present in more than 60% of the accessions (Fig. 6i). Five alleles, mainly clustering together, were classified as HR-associated, with allele34 being the most represented.

## Discussion

The ever-changing pathogen landscape imposes strong selective pressure on plants, leading to a fast evolution and extensive genetic diversity of NLRs, both at the inter- and intra-specific levels^1,4,51^. Pan-NLRome studies in various crop species^9–12,17,19^ highlighted that exploring SVs, CNVs, and NLR allelic diversity within species is essential for a complete immunity characterization. The pan-NLRome that we produced using 143 melon accessions from diverse botanical groups and global geographical origins represents the most extensive effort to date for NLR diversity characterization in this crop, and one of the most comprehensive across crop species. Moreover, we proposed original GWAS methods to untangle the associations between specific NLR genes/alleles and resistance to pathogens.

Complete and accurate pan-NLRome reconstruction depends of at least four factors: the completeness of accessions sampling, the quality of the genome assemblies, the gene models prediction accuracy, and the fidelity of NLR annotation. The 143 accessions presented diverse resistance patterns to several pest and pathogens and constituted the largest NLR diversity studies to date for any crop species using long reads. We used NAS^33^ to produce long reads from their NLR-containing genome regions, resulting in highly accurate assemblies. Moreover, we developed a pipeline for gene prediction and exhaustive NLR annotation. Unlike RenSeq, NAS captures entire NLR regions rather than isolated genes. Moreover, NAS improves contig anchoring to reference genomes and reveals the full genomic context of NLR regions. Notably, with NLRome-GWAS, we discovered candidate non-NLR genes within NLR region Chr05.3, statistically associated with resistance to Fusarium wilt race 1. This highlights the ability of our method to capture non-NLR resistance genes clustered with NLRs by shared evolutionary processes^52^.

The NLR content ranged from 77 to 104 per accession, principally due to expansion and contraction of clusters, with nearly all the NLR clusters existing in all accessions. This number represents a reduced NLR repertory compared to crops with a similar or smaller genome size^9,10,20^, but follows the same pattern found in other species from the *Cucurbitaceae* family^53,54^. Moreover, classification of annotated NLRs revealed a near 1:1 TNL/CNL ratio in melon, similar to that observed in some other *Cucurbitaceae* species^54–56^, but contrasting with the CNL-dominated profiles of other dicot families like *Solanaceae*^57^, and the TNL-dominated profile of *Brassicaceae*^58^. The RNL number was low and conserved across melon accessions, aligning with the results obtained in other crop species from diverse families^59^. RNLs often act as signaling helpers and do not directly participate in pathogen recognition^59^, which may explain their low number and reduced CNV. Additionally, we observed that RNLs were not mixed with other NLR classes in the melon genome, further supporting their role as helper NLRs that may require distinct regulation and broad availability to assist multiple sensor NLRs.

NLR expansion is thought to be constrained by a biological cost, including the energy required for transcription, translation, and regulation of expression^60^, which may explain the low NLR content in *Cucurbitaceae*. However, how species within this family cope with high pathogen pressure despite their reduced NLR repertory has remained unclear. Analysis of NLR diversity through orthogroups displayed that less than 100 accessions were needed to reach 95% of NLR saturation. Nevertheless, some orthogroups can gather hundreds of allelic variants or paralogs conferring resistance or susceptibility to diverse pathogens^39,41,61^. In this study, the 167 *Vat* allelic variants were grouped in orthogroup 3, some of them already reported to confer resistance to *Aphis gossypii* and powdery mildew^43^. Similarly, *Prv* and *Fom-1* alleles, involved in response to Fusarium wilt races 0 and 2, and to papaya ring-spot virus, respectively^40^, were also clustered together. This suggests that the usual analysis of NLR diversity via orthogroups^10,11,20^ hides the real existing allelic diversity, producing a tunnel vision for the association of NLR alleles and pathogen effectors. This underrepresentation of diversity is further supported by the different saturation patterns observed between our sequence-based pan-NLRome graph (unsaturated) and the orthogroup-based pan-NLRome. We annotated 13,323 NLRs that were clustered into 3,623 nr NLR alleles, and alleles’ saturation analysis showed a trend far from reaching a plateau after the analysis of 143 accessions. Potential misannotations of intron–exon structures, such as the one we identified for the *Fom-1* gene, are unlikely to have inflated the reported allelic diversity. After reannotation of *Fom-1* in 127 accessions, we still identified 34 nr alleles, yielding a comparable diversity rate. Likewise, manual curation of *Vat* alleles resulted in 167 nr sequences out of 726 annotated alleles, supporting the reliability of our diversity estimates. To further investigate allelic richness and account for NLR alleles that might exist in the species but were not detected due to sample size limitation, we estimated different measures of alpha diversity, a term already used for genetic diversity characterization^62–64^. Asymptotic Chao2 estimators suggested that almost half of the NLR alleles existing in the species remained undiscovered. Species may cope with pathogens’ pressure by enlarging their NLR repertory in a single genotype (increasing the number of NLR orthogroups) or population manner (increasing the allelic diversity within orthogroups). The here reported extreme NLR allelic diversity, never explored at this level, could be a reason explaining the limited number of NLR loci in melon, aside of the reported expansion of cell surface receptors in *Cucurbitaceae*^56^.

NAS on NLR regions also allowed the development of GWAS and post-GWAS methods to pinpoint specific NLR resistance associations that cannot be found with classical single-reference SNP-based GWAS. We proposed two targeted-NLR GWAS: one using a pan-NLRome graph as a reference, firstly applied in this context to our knowledge; and a second, reference-free k-mer-based method, previously applied to WGS^30,32^ but limited by high computational demands and the lack of multiple assemblies to map significant k-mers absent from the reference. To our knowledge, only ref. ^65^ applied k-mer-based GWAS to targeted NLRs, though lacking a broad NLR’s genomic context, and thus missing R genes clustered with NLRs. In our study, graph- and k-mer-based GWAS showed higher accuracy than SNP-based GWAS in identifying the previously cloned *Fom-1* and *Fom-2* alleles. Using a relatively small diversity panel, the k-mer based method presented an extreme precision, with the top k-mers mapping exactly to both genes. This method, leveraging a portfolio of NLR regions to locate significant k-mers absent from a single reference, is likely the most suitable as it does not require complex graphs construction or high-quality assemblies to map back significant k-mers. By combining the estimation of sampling completeness with discovered R allelic diversity, we predicted the number of *Fom-2* and *Fom-1* alleles still undiscovered. For Fusarium races 1 and 2, we observed four and seven resistance alleles with estimated sample completeness of 85 and 80%, suggesting that only one or two R alleles remain to be found for these genes. *Agrestis*, *flexuosus* and *acidulus* groups showed the highest Chao2 NLR allelic diversity estimators, suggesting that remaining NLR diversity should be primarily explored there.

By combining extensive diversity with innovative GWAS and accurate NAS assemblies, we identified alleles that can be arranged within haplotypes, a process likely portable to other species. By extending phenotyping to many pests and pathogens, haplotypes could be associated to multi-resistances. As some NLRs are also likely responsible of tolerance to abiotic stresses^66^, extending phenotyping to these stresses could lead to a breakdown in breeding approaches by selecting haplotypes rich in alleles responsive to multiple stresses while excluding high susceptibility ones. This would lead to the creation of varieties highly adapted to the current agroecological challenges.

## Materials and methods

### PLANT MATERIALS AND GENOME SEQUENCING

#### Samples selection

We selected 151 melon accessions belonging to 17 botanical groups and originated in very diverse areas of the world (Supplementary Table 1; Extended Data Fig. 1). We chose these accessions due to their resistance to different pests and pathogens affecting the Mediterranean basin, based on historic phenotyping data recorded at INRAE GAFL, Avignon, France. We recovered the seeds from the INRAE Center for Vegetable Germplasm in Avignon, France ^67^ and cultivated them under greenhouse conditions at INRAE GAFL. We sampled young plant leaves and froze them in liquid nitrogen for subsequent DNA extraction.

#### DNA extraction, libraries preparation and sequencing

We extracted genomic DNA and we conducted Oxford Nanopore Technologies (ONT) long-reads library preparation and NAS sequencing as described in ref. ^33^, using the Native Barcoding Kit 24 V14 (SQK-NBD114.24) (ONT, Oxford, UK). To determine the acceptance or rejection of reads, we provided 15 NBS-containing regions of ANSO-77 (Supplementary Table 12), masked for repetitive elements as presented in ref. ^33^, to the MinKNOW software (ONT, Oxford, UK). Using complementary NLR annotation tools like NLRAnnotator v. 2.1b^68^, and analyzing the NLR set of previously released *Cucumis melo* assemblies^69–72^, we discovered that six isolated NLR genes or domains were missed in the reference used for NAS. These corresponded to a TIR and TIR-NBS domains in chromosome 6, a NBS and TIR domains in chromosome 9, a CC-NBS in chromosome 10, and a TIR domain in chromosome 12. We included these six isolated NLRs with a flanking 20kb buffer zone into the provided reference for NAS (Supplementary Table 12) only for the last sequencing run, used to sequence the accessions Tibish 951, GAUCHO ISLA, Cum 412, HSD 195, WM 7 and Charentais Lavergne.

Accessions ANSO-77, DOUBLON, CHANG-BOUGI, CANTON, ANANAS, VEDRANTAIS, PI 414723 and ZHIMALI were sequenced in refs. ^33,73^. For the rest of accessions, we performed NAS on whole R10.4.1 PromethION flowcells as described in ref. ^33^ for CHANG-BOUGI. Barcodes and multiplexing configuration for each sequencing run are specified in Supplementary Table 13. Due to occasional barcode imbalances in the sequencing pool, some of these accessions were sequenced across multiple runs to achieve sufficient coverage for assembly (Supplementary Table 13). We extended runs during 120 hours, and we performed a library reloading (washing flush) when the percentage of sequencing pores dropped to 10-15%. Due to regular ONT softwares updates, we used five different versions of MinKNOW for the sequencing of the accessions (Supplementary Table 13). Raw ONT FAST5 files were live base-called during the PromethION run with Guppy (ONT, Oxford, UK) in “super accurate base-calling” mode. We used four different versions of Guppy due to regular software updates.

We sequenced with short reads all accessions except *Tibish 951, Cum 412, HSD 195,* and *Charentais Lavergne.* We used short-reads 150 bp paired-end sequencing technologies from Illumina NovaSeq6000 (San Diego, CA, USA) and/or MGI DNBSEQ-G400RS with HotMPS High-throughput Sequencing chemistry (MGI, Shenzhen, China) (Supplementary Table 13), ensuring a minimum average depth of 15×. Libraries preparation and sequencing followed the protocols described in ref. ^73^.

### GENOME ASSEMBLIES, EVALUATION AND ANNOTATION

#### NAS reads processing and targeted genome assembly

For the accessions sequenced here, reads were processed as outlined in ref. ^33^ for CHANG- BOUGI. We filtered out reads shorter than 5 kb prior assembly. We used a defined procedure to assemble the targeted NLR regions from NAS reads. First, we performed assemblies using SMARTdenovo v. 2018.2.19^74^ as we previously demonstrated its high performance, accuracy and contiguity for the assembly of NLR clusters in melon^33^. We filtered SMARTdenovo assemblies as described in ref. ^33^. We assessed the accuracy and completeness of the assembled contigs by aligning the reads back to them using Minimap2 v. 2.28^75^ with parameters ‘-ax map-ont -r2k’ and conducting visual inspection of the mappings using IGV v. 2.17.1^76^. Alignments presenting polymorphisms were mainly due to miss-assemblies of segment duplications, missing clusters, or high heterozygosity rates, as current ONT-based assemblers typically collapse both haplotypes^77^. We retrieved contigs presenting a certain level of polymorphisms, and we reassembled them with other ONT-based assemblers: Canu v. 2.2^78^, Shasta v. 0.11.1^79^, NextDenovo v. 2.5.0^80^ and Flye v. 2.9.5^81^. We used default parameters and added options ‘genomesize = 7m –corrected –trimmed –nanopore’ for Canu, ‘--config Nanopore-R10-Fast- Nov2022 --MinHash.minHashIterationCount 100’ for Shasta, ‘genome_size=7m’ for NextDenovo, and ‘--nano-hq -g 6.5m --read-error 0.03’ for Flye. After this step, some NLR regions were still showing polymorphisms in a near to 50% rate, indicating collapsed assemblies due to the presence of a certain degree of heterozygosity. In a third step, we used a HERRO^82^ deep-learning correction + Hifiasm v. 0.19.9^83^ procedure to achieve partially phased ONT-only assemblies of problematic regions. We ran HERRO using the model ‘model_R10_v0.1.pt’. We provided HERRO-corrected NAS reads to Hifiasm as Hifi reads, and the original NAS reads as ONT reads, and we used parameters ‘-l3 --hg-size 7.5m --purge-max 200 -s 0.25’.

We successfully recovered the six NLR genes or domains not included in the reference used for NAS along with 20 kb of flanking genomic sequence through a reference-guided assembly process. We provided raw NAS reads along with WGS short reads selected for each missed NLR to the software Geneious Prime v. 2022.1.1 (Biomatters, Auckland, New Zealand), using the ANSO-77 assembly^33^ as reference.

After all the steps, we merged all assembled NLR regions (15 NAS-targeted and six reference-guided) for each of the accessions. We aligned assembled NLR regions to the 21 NLR regions of ANSO-77 in order to get their genomic positions. We used Minimap2 with default parameters and adding ‘-f 0.02’ for contigs alignment.

#### Quality and contiguity assessment of *de novo* assemblies

We mapped the reads used for assembly back to the NAS-targeted assemblies to evaluate their accuracy and completeness, using Minimap2 with parameters ‘-ax map-ont -r2k’. To prevent noisy reads originating from non-target regions from mapping to the target assemblies, we constructed a chimeric genome for each accession. These chimeric genomes were constituted by the target NLR regions for each accession and the non-target genome of ANSO-77. We called SNPs from the mapping files using htsbox pileup v. r346^84^ with parameters ‘-S 1000 -q 60 -Q 5 -s 7’. Following the method proposed in ref. ^10^, we generated a measure of assembly quality (accuracy and completeness) for each assembled contig by comparing the number of SNP calls (nb_snp) to the total size of the contig (total_nt), as expressed in equation (1):

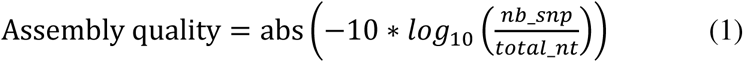

We measured contiguity for each NLR region and accession as the ratio of the number of haplotypes assembled for that region to the total number of assembled contigs.

#### Automatic and manual gene prediction

We performed automatic structural gene annotations on the assembled regions with Helixer v. 0.3.3^85^, as we previously reported its high accuracy to annotate well known NLRs^33^. We complemented this annotation with an expert manual annotation of *Vat* genes between the M5 and M4 markers^44^. We combined both annotations using custom Bash scripts.

### NLRs ANNOTATION AND ANALYSIS OF THEIR GENOMIC DISTRIBUTION

#### NLR identification, classification and counting

We used four different tools to identify and classify NLRs: NLGenomeSweeper, NLRAnnotator v. 2.1b, RGAugury v. 2.0^86^ and Resistify v. 0.5.1^14^. These tools complement each other as they use different evidences to identify NLRs. NLGenomeSweeper and NLR-annotator use nucleotide sequences as input, while RGAugury and Resistify are protein-based. We developed a Bash script that combined the four annotations to generate a consensus. We lifted the annotations from NLGenomeSweeper and NLR-Annotator on the structural gene annotation from Helixer. We discarded nucleotide-based annotations not supported by any protein-based annotator, though these were usually very few. We selected the NLR domain composition based on majority vote method. When no NLR classification was predominant, we prioritized Resistify, followed by RGAugury, NLRAnnotator, and NLGenomeSweeper. Resistify was preferred as it integrates multiple approaches and leverages deep learning^14^.

Similar to ref. ^10^, we considered as NLR genes those that contained at least an NBS, TIR or a RPW8 domain, excluding isolated LRR or CC domains. Based on the different domain combinations, we classified NLRs into 12 subclasses: N, NL, TN, TNL, CN, CNL, RN, RNL, TIR, RPW8, TX and OTHER. For convenience, C, N, L, T, R and X abbreviates CC, NB-ARC, LRR, TIR, RPW8 and unknown domains. Genes classified as OTHER exhibited chimeric domain architectures. We grouped these subclasses into five classes: TNL, CNL, RNL, NL, and OTHER, based on the presence of the N-terminal domains TIR, CC, and RPW8.

We counted NLRs for each NLR region and accession. For regions assembled into two pseudo-haplotypes, produced by Hifiasm, we estimated both the minimum and maximum possible NLR counts (Extended Data Fig. 9).

### NLROME DIVERSITY CHARACTERIZATION

#### Orthogroups identification and allele-based NLRome construction

We constructed a pan-NLRome based on orthogroups. We used Orthofinder v. 3.0^87^ with default parameters to cluster all annotated NLR alleles across the 143 accessions into orthogroups. In parallel, we constructed an allele-based pan-NLRome using a protein-clustering approach. For each NLR region, we clustered NLR alleles sharing identical protein sequences, constituting nr NLR alleles. We generated P/A matrices of orthogroups and nr NLR alleles using custom Bash scripts.

Based on the frequency of orthogroups and nr NLR alleles observed across the 143 accessions, we classified them into three different pan-genome categories: core (present in at least 90% of accessions), shell (present in more than 10% and less than 90% of accessions), and cloud (present in 10% of accessions or less).

#### Saturation analysis

We generated growth saturation curves from the P/A matrices, by randomly selecting an increasing number of assembled accessions (from one to 143). We performed 1,000 iterations at each specific number of accessions. The number of orthogroups and the number of nr NLR alleles were identified for each of the replicates and averaged, obtaining also a standard deviation. We used custom R scripts for constructing and plotting the growth saturation curves.

#### NLR allelic richness calculation

We applied four metrics to assess NLR allelic diversity at the protein level. Allelic richness provided a standardized measure of diversity by normalizing the observed number of nr NLR alleles by the average NLR count in each region^88^. To further investigate allelic richness and account for alleles that might exist but were not detected due to sample size limitation, we calculated different measures of alpha diversity, a concept already applied to study genetic diversity^62–64^. We measured three indices of alpha diversity: the Chao2 index^89^, an estimator of richness that corrects for alleles that might be present but not detected due to insufficient sampling; the Shannon index, that balances richness with evenness; and the Simpson index, similar to the Shannon index, but giving more weight to abundant nr NLR alleles^90^. We calculated these three indices in R with the package iNEXT^46^, using the functions ChaoRichness(), ChaoShannon() and ChaoSimpson() with 95% confidence level for intervals. We also calculated Chao2 richness among botanical groups and geographical origins by rarefaction to the smallest sample size, reducing biases coming from unequal number of samples^45^. We considered groups containing at least seven accessions. We used the function estimateD() from the iNEXT package, with parameters ‘q=0 conf=0.95 base=“size” datatype=“incidence_raw”’.

To assess the representativeness of our accession sampling among NLR regions, we constructed sample completeness curves using the rarefaction and extrapolation framework implemented in iNEXT. We used the iNEXT() function, setting 500 bootstrap replicates, confidence level intervals of 95%, and 80 knots (equally-spaced sample sizes for which diversity estimate was calculated).

### PRE-GWAS ANALYSES

#### Symptoms severity scoring and phenotypes analysis

We evaluated the phenotypic response of the selected panel of accessions to Fusarium wilt (*Fusarium oxysporum* f. sp. *melonis)* race 1 MATREF isolate, Fusarium wilt race 2 R ISF isolate, and ZYMV strain E9 pathotype 1. For Fusarium races 1 and 2, we conducted two independent trials, each consisting of two randomized complete blocks. Additionally, a third randomized incomplete block was included in one of the trials for Fusarium race 2. We used trays containing 80 plants and 16 controls (arranged in 12 × 8) for one trial, and trays of 48 plants (6 × 8) for the other. Trays were filled with potting soil treated with Previcur® (Bayer AG, Leverkusen, Germany) to prevent *Pythium* contamination. We sowed 12 seeds per genotype, block and trial. We grew plants in a climate-controlled chamber under a 16/8-hour light/dark cycle at 25 °C during the day and 20 °C at night. We performed inoculation at the expanded cotyledon stage, when the first true leaf was appearing (7 to 10 days after sowing). Each tray was dipped into 1 L of spore suspension at a concentration of 10⁶ spores/mL, until complete absorption by capillarity. The symptoms severity was scored for each plant 18 days after inoculation, when resistant and susceptible checks revealed the expected symptoms. We used the following notation scale: 0 – no symptoms of yellowing and wilting, 1 – light symptoms of yellowing and wilting, 2 – yellowing, wilting and necrosis, stunting, 4 – dead plant. We removed plants with late germination or *Pythium* symptoms before scoring. From the recorded phenotypic data, we considered as outliers and discarded all scores higher or lower than two standard deviations within each accession. Afterwards, we manually deleted blocks with less than five plants per genotype, and genotypes phenotyped in less than two blocks. We estimated the contribution of the experimental effects to the recorded phenotypes using the mixed linear model (MLM) presented in equation (2),

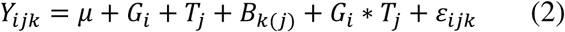

where Y_ijk_ represents raw phenotypic observations; G_i_ is the random effect of genotype i, with distribution 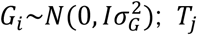 the random effect of trial j, assumed to be distributed as 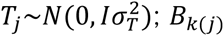 the random effect of the block k nested within the trial j, assumed to be distributed as 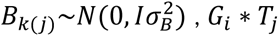 the interaction between G_i_ and T_j_, and ε_ijk_ the residual error effect, expected to follow distribution 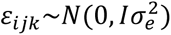. We computed this mixed model with the lmer() function from the R lme4 package^91^.

We estimated heritability using the variance of the effects from the previous model, as described in ref. ^92^, following equation (3),

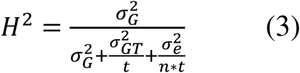

where 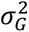 and 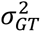 are the variance components associated to the genotypes and genotypes-by-trial interaction, respectively; t the number of trials (two); n the average number of observations per genotype and trial over the whole design; and 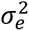 the residual variance.

We condensed up Fusarium wilt observations to single Best-Linear-Unbiased-Predictors (BLUPs) for each genotype for further GWAS. As symptoms severity scores for both Fusarium wilt phenotypes were categorical, we used a generalized Poisson mixed linear model presented in equation (4) to extract BLUPs,

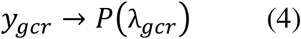

where y_gcr_ represents the symptoms severity scores following a Poisson law of parameter λ_gcr_, with log(λ_gcr_) = Wv + Gg, where v is the vector of fixed effects (trial); W is the fixed effects incidence matrix; g is the vector of random effects (genotype, block and interaction genotype-by-trial), assumed to follow a distribution 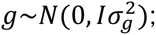 and G is the random effect incidence matrix. We used the glmer function from R lme4 package to build the model. We extracted BLUPs from the model using the ranef() function from the same package.

For ZYMV, we implemented five incomplete block designs, each testing a different subset of accessions. We conducted experiments in insect-proof greenhouses using plastic trays containing 60 plants arranged in a 10 × 6 grid. Trays were filled with potting soil, and we sowed ten seeds per genotype in single rows. We grew plants under controlled conditions until they reached the cotyledon stage, approximately 7–10 days after sowing. At this point, we performed a mechanical inoculation using an extract prepared from fresh ZYMV-infected zucchini leaves as described in ref. ^93^. We used a binary scoring system: 0 – no symptoms or flaccid necrosis; 1 – flaccid necrosis, dead plant. Accessions with fewer than five individual phenotypic observations were excluded from downstream analyses. We observed no phenotypic variation within any accession. Therefore, we used this binary phenotype directly in GWAS.

#### SNP matrix construction

We used a SNP matrix from short-reads data of 226 accessions available at INRAE-GAFL (Supplementary Table 11), including 147 presented in this study. ANSO-77 was used as reference assembly^33^ for matrix construction. The whole process was applied using a Snakemake pipeline (https://forgemia.inra.fr/gafl/pipelines_snakemake/wgs_gatk). This matrix was filtered by retaining only biallelic variants. We removed variants having missing data or genotype quality (GQ) < 20 or read depth (DP) < 7 in more than 3% of the samples, as well as those with a heterozygosity rate exceeding 50%. Then, we set as missing variants those presenting GQ below 15 or DP below 5. This filtering process resulted in a high-quality variant matrix, hereafter referred to as MA_NLRome_, that included 3.84 million SNP calls.

#### Genetic structure and kinship calculation

We explored population structure groups using the MA_NLRome_ matrix. We selected 8,000 independent SNPs from this dataset by doing LD pruning with Plink v. 1.90^94^, in order to optimize computational efficiency. We used a sliding window size of 100 kb, a step size of one SNP, and a squared correlation coefficient (r^2^) threshold of 0.2. We calculated population structure via STRUCTURE v. 2.3.4^95^, including 1-10 possible groups and 10 runs per group. Each run consisted on 250,000 burn-in steps and 500,000 Markov chain Monte Carlo (MCMC). We collected STRUCTURE results and built Evanno plots^96^ for grouping decisions using the STRUCTURE Harvester^97^.

We selected phenotyped accessions from MA_NLRome_ and generated kinship matrices for each subset by applying equation (5) using the A.mat() from the R package rrBLUP^98^:

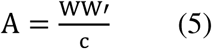

where W_ik_ = X_ik_ + 1 − 2p_k_, being i the individual and k the SNP, X the genotype matrix, and p_k_ the frequency of the allele 1 at SNP k; and c = 2 ∑_k_ p_k_ (1 − p_k_).

#### Pan-NLRome graph construction, variant extraction, and nodes-based P/A matrix construction

We constructed a pan-NLRome graph of the 21 NLR regions across the 143 assembled accessions using PGGB^27^. Each of the 21 NLR regions was constructed as a sub-graph, containing 154, 150, 153, 150, 161, 146, 148, 162, 151, 143, 143, 153, 153, 149, 153, 153, 143, 143, 143, 156 and 143 assembled haplotypes, respectively. As we sequenced NLR clusters using NAS, the flanking regions surrounding each cluster varied in length between accessions. To ensure consistency and avoid including these additional sequences in the statistical analyses extracted from the NLRome graph, we trimmed “dangling” ends prior to NLRome graph construction. We used Minimap2 with parameters ‘-ax map-ont -r2k’ to align each assembled region to the ANSO-77 reference for NAS, and we retained the matching part for downstream analyses. We independently used the tools wfmash v. 0.10.5 with parameters ‘–s 500 –l 2,500 –p 95 -k 19 -H 0.001’ for all-to-all alignment of input sequences; seqwish v. 0.7.11 with parameters ‘-B 10000000 -k 19 -f 0’ for graph induction; and GFAffix v. 0.2.0 with default parameters for graph normalization. We produced NLRome growth histograms and other statistics using panacus v. 0.2.3^42^ with default parameters and including the path of ANSO-77 as reference. We ran all these pan-genome tools through Pan1c (https://forge.inrae.fr/genotoul-bioinfo/Pan1c/pan1c/-/tree/main), a Snakemake workflow for creating pan-genomes at chromosomic scale. We extracted variants from the graph using vg deconstruct v. 1.63.1^99^ with default parameters and including ‘-a’ to report all bubbles, including nested ones.

We constructed a P/A matrix of graph nodes across accessions in the pan-NLRome graph, containing 949.68K variant calls. This matrix was generated from the GFA graph file using a custom Python script and was subsequently filtered for each phenotype subset and used for GWAS (see GWAS analyses section). We calculated kinship matrices from the filtered P/A matrices of nodes across accessions using the R function A.mat(), described in equation (5).

#### K-mers extraction and k-mer-based P/A matrix construction

We developed a Snakemake pipeline that makes use of the approach developed by ref. ^30^ to generate a k-mer P/A matrix across accessions (Extended Data Fig. 10). We specified a k-mer length of 31, low and top thresholds for k-mer count in an accession of 8 and 400, MAF of 0.04 and two minimum percentage of appearance in each strand form of a k-mer. The pipeline used KMC v.3.2.4^100^ to count canonized and not canonized 31-mers. Then, it uses the kmersGWAS v. 0.3^30^ library to create a consensus of k-mers counts per accession, convert the list of k-mers to a matrix of P/A, and generate a kinship matrix from the k-mers count. We restricted k-mer counting to NLR regions. We provided NAS reads for the 15 targeted regions, and a combination of raw NAS and short reads for the six additional regions containing isolated NLRs. The generated matrix contained 20.24 million k-mers. This matrix was filtered for each phenotyped subset prior to GWAS analyses (see GWAS Analyses section).

### GWAS ANALYSES

We performed GWAS using three different genotype matrices: a SNP matrix obtained from short-reads mapping and variant calling, a matrix of P/A of nodes from the pan-NLRome graph, and a matrix of P/A of k-mers within the defined NLR regions. For each phenotype, we used the same set of accessions and applied the same GWAS model across SNP-, graph- and k-mer-GWAS to ensure comparability of results. Prior to GWAS analyses, the three matrices were sample-filtered by selecting common samples, and variant-filtered using a MAF of 0.06 (representing 7-9 accessions).

To perform GWAS on both Fusarium wilt phenotypes, we used the multi-locus mixed model (MLMM) implemented in the R package mlmm^101^. This model follows a multistep approach. It first applies a null mixed linear model (MLM) and then iteratively includes the top SNP from the previous step as a fixed effect in subsequent steps. We performed the multistep approach for SNP-based GWAS, but we only performed the first step (before including top variants as cofactors) for k-mer- and node-based GWAS. We defined the last MLMM step when no further SNPs were significant or when there was no longer phenotypic variance under genetic control. To perform GWAS on the ZYMV flaccid necrosis phenotype, we used a binomial GLMM, more adapted to binary phenotypic data. We used the GMMAT package in R^102^, with the functions glmmkin() to fit the binomial GLMM and glmm.score() to perform GWAS.

We evaluated four models for SNP-based GWAS: null (no kinship or population structure correction), kinship-only (K), population structure with five genetic groups (Q5), and combined model incorporating both kinship and structure (KQ5). For P/A matrices derived from graph nodes and k-mers, we tested only the null and kinship models. We selected the model that best fitted the data via Q-Q plots generated with the qqman R package^103^. Equation (6) provides a general formulation that encompasses all tested model combinations and applies to both MLMM and GLMM frameworks,

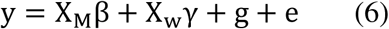

where y is the phenotype vector (BLUPs for Fusarium wilt data or raw phenotypes for ZYMV); β is the vector of allelic effects; X_M_ is the matrix with genotype values for M accessions at the 560 to 618K SNPs being tested; γ is the fixed effects vector, including population genetic structure and/or top SNP (for MLMM), when appropriate according to the applied model; X_w_ is the fixed effects incidence matrix for W effects; g is the random genotype effect vector that follows the distribution 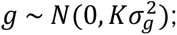 K is the kinship matrix; 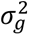 is the genetic variance, e is the error vector that follows the distribution 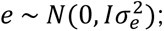 and 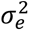 is the residual variance. g and e are assumed to be independent.

We defined a Bonferroni significance threshold with a genome-wide error risk of α=0.05 corrected by the number of independent variants in all cases. In SNP-based GWAS, the number of independent SNPs was determined with Gao’s method^104^ using a 100 kb sliding window. In k-mer GWAS, we approximated the number of independent k-mers as the count of distinct P/A patterns observed across accessions, assuming that k-mers tagging dependent variants will typically show the same P/A pattern. In the GWAS with graph nodes, we approximated the number of independent nodes as the count of top-level snarls, assuming that nested nodes are dependent of the top-level ones.

### POST-GWAS ANALYSES

#### Local linkage disequilibrium for QTL detection

We defined QTLs from SNP-based GWAS results by calculating LD around the top SNPs. We selected the top SNP and measured the r² of pairwise LD 10 Mb around it, using the Measure.R2VS function from the LDcorSV R package^105^, and including population structure and kinship in the model. From these r² results, we defined the QTL around the selected top SNP using a sliding window approach, iteratively expanding left and right boundaries by 100 kb around the SNP, until we did not find SNPs over a 0.2 r² threshold in either direction. We repeated this procedure for the subsequent significant SNPs located outside the previously defined QTL regions, until no significant SNPs remained outside defined QTLs.

#### Anchoring of top-associated k-mers

We mapped top-associated k-mers identified through GWAS to the assembled NLR regions of the accessions in which they were present, using a custom Bash script (Extended Data Fig. 10). First, we selected the k-mers presenting a -log of p-value over the corrected α threshold, and we identified the accessions that presented those k-mers (from the P/A matrix). Then, we clustered k-mers presenting exactly the same p-value, assumed to evidence the same structural variant, and we did a local assembly of each k-mer block using minia v. 3.2.6^106^ with parameters ‘-kmer-size 15 -abundance-min 1 -no-tip-removal’. Afterwards, we aligned the assembled k-mers to the assemblies of NLR regions of the accessions where they were present using BLAST v. 2.12, with parameters ‘-word_size 7 -outfmt "6 std qlen" -max_target_seqs 5’. We retained only BLAST hits with 100% identity to the top-associated k-mers. Finally, we added the feature annotations of the k-mer positions on the assemblies with bedtools v. 2.29.2^107^.

#### Anchoring of top-associated nodes

We recovered the position of top-associated nodes identified through GWAS on the assembled NLR regions of all analyzed accessions. First, we determined the positions of top-associated nodes along accession-specific paths in the graph using the odgi position command included in PGGB v. 0.5.1. Subsequently, we extracted these accession paths in FASTA format using the vg paths command from the same version of PGGB. The sequences of these paths differed from the original ones, as we trimmed “dangling” ends longer than the ANSO-77 reference for graph construction. We used Minimap2 with parameter -f 0.02 to align the extracted graph paths to the original annotated sequences and establish correspondences between them. Based on these alignments, we lifted the coordinates of the top-associated nodes from the graph paths to their corresponding positions in the original annotated sequences.

#### Manhattan plots generation

We generated Manhattan plots using the whole-genome assembly of ANSO-77 as a reference, incorporating all SNPs, as well as significantly associated k-mers and graph nodes. We built these plots using the R ggplot2 package^38^. We plotted all significantly associated k-mers, regardless of whether they aligned with 100% identity to the assembly. We determined their positions as described above, but in this case, we retained hits with the best E-value rather than restricting to perfect matches. Similarly, we plotted all significantly associated nodes, even when the path of ANSO-77 was not contained in that node. We used this approach to illustrate that relying on mapping on a single reference, a common practice in k-mer-based GWAS, can lead to misleading or erroneous localization of associations.

We generated zoomed Manhattan plots spanning 50 kb upstream and downstream of the midpoint of the top-associated gene, as identified by SNP-, k-mer- and/or node-based GWAS. We used the NLR assemblies of ANSO-77 and Isabelle as references, as these two accessions consistently exhibited contrasting phenotypes for the three studied diseases. In this case, we only included top-associated k-mers and graph nodes that were present in ANSO-77 and Isabelle.

#### Alleles identification and phylogenetic trees generation

We extracted alleles of top-associated genes across all assembled accessions for the three studied phenotypes using a custom Bash script. First, we aligned the top-associated gene to each phenotyped accession using Minimap2 v. 2.24 with the parameter –f 0.02. We used as reference the top-associated genes ME00791#1#utg22_Chr11.1_000033 (gene *Fom-2* from Isabelle) for Fusarium wilt race 1; *Fom-1* gene (splice variant A) from MR-1 for Fusarium wilt race 2, retrieved from GenBank accession JX295632.2; and ME00509#1#utg45_Chr05.3_000021 from ANSO-77 for ZYMV-induced flaccid necrosis. Then, from each accession, we retrieved gene annotation corresponding to the aligned region using the command intersect from bedtools v. 2.29, we extracted the protein sequences. Multi-alignments of those proteins revealed erroneous Helixer intron-exon annotations in 12 *Fom-2* alleles and all *Fom-1* alleles. Therefore, we used Liftoff v1.6.3^108^ with default parameters and using same gene references to retrieve the correct annotations. We confirmed the accuracy of new annotations by protein multi-alignments.

We phylogenetically analyzed the extracted protein alleles to assess their evolutionary relationships. We performed alignments using MAFFT v. 7.490^109^ with default parameters. We fed IQ-TREE v. 1.6.12^110^ with these alignments, using parameters ‘-bb 10,000 -alrt 10,000’, to generate maximum likelihood phylogenetic trees. The best-fit substitution model was selected automatically by ModelFinder, embedded in IQ-TREE, using the Bayesian Information Criterion (BIC) for decision. The models selected were HIVb+F+I for *Fom-2*, HIVb+R2 for *Fom-1* and JTTDCMut+F+R2 for the first candidate gene of ZYMV-induced flaccid necrosis. We used the online version of iTOL v. 7^111^ to display and annotate the phylogenetic trees generated with IQ-TREE.

### DATA ANALYSIS AND FIGURE GENERATION

We used Bash, Python and R scripts to analyze data and prepare datasets. We generated figures and plots using RStudio v. 2023.09.1, and we used Inkscape v. 0.91 to manage and refine SVG outputs, as well as assemble multi-panel figures. We generated dot plots between assemblies using D-GENIES^112^.

## Data availability

ONT-NAS raw sequencing data is available at the NCBI database under BioProject PRJNA1127998. The large sequencing_summary.txt files used for read filtering by end reason prior assembly are available from the corresponding author on reasonable request. Short-reads raw sequencing data is also available at NCBI under BioProjects PRJNA1164662 (for CANTON), PRJNA662717 (for ANSO-77), PRJNA662721 (for DOUBLON), PRJNA1164664 (for ZHIMALI), PRJNA1164667 (for PI 414723), PRJNA1164698 (for VEDRANTAIS), PRJNA1164660 (for ANANAS), and PRJNA1273021 (for all the rest of accessions). The targeted assemblies of 21 NLR regions across the 143 accessions, together with gene annotation and NLR annotation is available at the Recherche Data Gouv database under ref. ^113^. The *Fom-1* and *Fom-2* reannotation is also provided at this database. Raw phenotyping data, as well as the pan-NLRome graph GFA file, the SNP matrix (MA_NLRome_), the k-mer P/A matrix, the graph nodes P/A matrix, and the k-mer kinship matrix are available in the Recherche Data Gouv database under ref. ^114^.

## Code availability

All scripts used for data analysis and plotting are accessible at the GitLab page indicated in the scripts_availability.txt file included in ref. ^113^.

## Acknowledgements

We are grateful to CEA-IbFJ-Genoscope for providing sequencing, bioinformatics and storage facilities. We thank the Genotoul bioinformatics platform Toulouse Occitanie (Bioinfo Genotoul, https://doi.org/10.15454/1.5572369328961167E12) for providing computing and storage resources; Sylvia del Valle from TAKII France SAS, as well as Daniel Bellon Doña and Maria Florencia Cocaliadis from BASF Nunhems for phenotyping the panel for resistance to Fusarium; and the INRAE Pangenome Working Group PANANNOT for valuable discussions and collaborative work.

## Authors’ contributions

J.B.M., A.B., and I.L. performed NAS sequencing and whole-genome short-reads sequencing. J.B.M generated NAS assemblies, produced gene and NLR annotations, performed NLR diversity analyses, developed and applied GWAS methods, and conducted all statistical analyses. P.F.R., N.B. and D.H. conceived the study and provided expertise. V.R.R. did DNA extraction and manually annotated the *Vat* cluster for all the assembled accessions. P.M. and K.L. managed seeds lots availability and contributed to disease phenotyping. J.L. provided bioinformatics expertise. J.B.M. and A.C. did the data submission. J.B.M., P.F.R., N.B. and D.H. wrote the manuscript. All authors read and approved the final manuscript.

## References

1. Jones, J. D. G. & Dangl, J. L. The plant immune system. Nature 444, 323–329 (2006).

2. Bentham, A., Burdett, H., Anderson, P. A., Williams, S. J. & Kobe, B. Animal NLRs provide structural insights into plant NLR function. Ann Bot 119, 827–702 (2017).

3. Duxbury, Z., Wu, C. & Ding, P. A Comparative Overview of the Intracellular Guardians of Plants and Animals: NLRs in Innate Immunity and Beyond. Annu. Rev. Plant Biol. 72, 155–184 (2021).

4. Barragan, A. C. & Weigel, D. Plant NLR diversity: the known unknowns of pan-NLRomes. The Plant Cell 33, 814–831 (2021).

5. Baggs, E., Dagdas, G. & Krasileva, K. NLR diversity, helpers and integrated domains: making sense of the NLR IDentity. Current Opinion in Plant Biology 38, 59–67 (2017).

6. Contreras, M. P., Lüdke, D., Pai, H., Toghani, A. & Kamoun, S. NLR receptors in plant immunity: making sense of the alphabet soup. EMBO reports 24, e57495 (2023).

7. Nishimura, M. T. et al. TIR-only protein RBA1 recognizes a pathogen effector to regulate cell death in Arabidopsis. Proceedings of the National Academy of Sciences 114, E2053–E2062 (2017).

8. Marchal, C. et al. BED-domain-containing immune receptors confer diverse resistance spectra to yellow rust. Nature Plants 4, 662–668 (2018).

9. Shang, L. et al. A super pan-genomic landscape of rice. Cell Res 32, 878–896 (2022).

10. Van de Weyer, A.-L. et al. A Species-Wide Inventory of NLR Genes and Alleles in *Arabidopsis thaliana*. Cell 178, 1260–1272.e14 (2019).

11. Ning, W. et al. The pan-NLRome analysis based on 23 genomes reveals the diversity of NLRs in *Brassica napus*. Mol Breeding 44, 2 (2024).

12. Thatcher, S. et al. The NLRomes of Zea mays NAM founder lines and *Zea luxurians* display presence–absence variation, integrated domain diversity, and mobility. Molecular Plant Pathology 24, 742–757 (2023).

13. Van Wersch, S. & Li, X. Stronger When Together: Clustering of Plant NLR Disease resistance Genes. Trends in Plant Science 24, 688–699 (2019).

14. Smith, M., Jones, J. T. & Hein, I. Resistify: A Novel NLR Classifier That Reveals Helitron-Associated NLR Expansion in Solanaceae. Bioinform Biol Insights 19, 11779322241308944 (2025).

15. Mo, C. et al. Complete genome assembly provides a high-quality skeleton for pan-NLRome construction in melon. The Plant Journal 118, 2249–2268 (2024).

16. Dolatabadian, A. et al. Characterization of disease resistance genes in the *Brassica napus* pangenome reveals significant structural variation. Plant Biotechnology Journal 18, 969–982 (2020).

17. Tang, D. et al. Genome evolution and diversity of wild and cultivated potatoes. Nature 606, 535– 541 (2022).

18. Walkowiak, S. et al. Multiple wheat genomes reveal global variation in modern breeding. Nature 588, 277–283 (2020).

19. Parada-Rojas, C. H. et al. A reference-quality NLRome for the hexaploid sweetpotato and diploid wild relatives. Preprint at 10.1101/2025.01.13.632774 (2025).

20. Liu, S. et al. The Diversity of NLRs in *Brassica rapa* Pan-genome. Preprint at 10.1101/2023.08.29.555307 (2023).

21. Lin, X. et al. Solanum americanum genome-assisted discovery of immune receptors that detect potato late blight pathogen effectors. Nat Genet 55, 1579–1588 (2023).

22. Seong, K., Seo, E., Witek, K., Li, M. & Staskawicz, B. Evolution of NLR resistance genes with noncanonical N-terminal domains in wild tomato species. New Phytologist 227, 1530–1543 (2020).

23. Gupta, P. K. Quantitative genetics: pan-genomes, SVs, and k-mers for GWAS. Trends in Genetics 37, 868–871 (2021).

24. Jayakodi, M., Schreiber, M., Stein, N. & Mascher, M. Building pan-genome infrastructures for crop plants and their use in association genetics. DNA Research 28, dsaa030 (2021).

25. Schreiber, M., Jayakodi, M., Stein, N. & Mascher, M. Plant pangenomes for crop improvement, biodiversity and evolution. Nat Rev Genet 25, 563–577 (2024).

26. Hickey, G. et al. Pangenome graph construction from genome alignments with Minigraph-Cactus. Nat Biotechnol 42, 663–673 (2024).

27. Garrison, E. et al. Building Pangenome Graphs. Preprint at http://biorxiv.org/lookup/doi/10.1101/2023.04.05.535718 (2023).

28. Vorbrugg, S., Bezrukov, I., Bao, Z., Xian, W. & Weigel, D. Gfa2bin enables graph-based GWAS by converting genome graphs to pan-genomic genotypes. Preprint at 10.1101/2024.12.05.626966 (2024).

29. Liu, Z. et al. Grapevine pangenome facilitates trait genetics and genomic breeding. Nat Genet 56, 2804–2814 (2024).

30. Voichek, Y. & Weigel, D. Identifying genetic variants underlying phenotypic variation in plants without complete genomes. Nat Genet 52, 534–540 (2020).

31. Karikari, B., Lemay, M.-A. & Belzile, F. k-mer-Based Genome-Wide Association Studies in Plants: Advances, Challenges, and Perspectives. Genes 14, 1439 (2023).

32. Jaegle, B. et al. k-mer-based GWAS in a wheat collection reveals novel and diverse sources of powdery mildew resistance. Genome Biology 26, 172 (2025).

33. Belinchon-Moreno, J. et al. Nanopore adaptive sampling to identify the NLR gene family in melon (*Cucumis melo* L.). BMC Genomics 26, 126 (2025).

34. Liao, W.-W. et al. A draft human pangenome reference. Nature 617, 312–324 (2023).

35. Marees, A. T. et al. A tutorial on conducting genome-wide association studies: Quality control and statistical analysis. International Journal of Methods in Psychiatric Research 27, e1608 (2018).

36. Jiang, Y. et al. Association Analysis and Meta-Analysis of Multi-Allelic Variants for Large-Scale Sequence Data. Genes 11, 586 (2020).

37. Lin, Y. et al. quarTeT: a telomere-to-telomere toolkit for gap-free genome assembly and centromeric repeat identification. Horticulture Research 10, uhad127 (2023).

38. Wickham, H. ggplot2. WIREs Computational Statistics 3, 180–185 (2011).

39. Ellis, J. G., Lawrence, G. J. & Dodds, P. N. Further analysis of gene-for-gene disease resistance specificity in flax. Molecular Plant Pathology 8, 103–109 (2007).

40. Brotman, Y. et al. Dual resistance of melon to *Fusarium oxysporum* races 0 and 2 and to Papaya ring-spot virus is controlled by a pair of head-to-head-oriented NB-LRR genes of unusual architecture. Molecular plant 6, 235–238 (2013).

41. Botella, M. A. et al. Three Genes of the Arabidopsis RPP1 Complex Resistance Locus Recognize Distinct *Peronospora parasitica* Avirulence Determinants. The Plant Cell 10, 1847–1860 (1998).

42. Parmigiani, L., Garrison, E., Stoye, J., Marschall, T. & Doerr, D. Panacus: fast and exact pangenome growth and core size estimation. Bioinformatics 40, btae720 (2024).

43. Boissot, N. et al. A highly diversified NLR cluster in melon contains homologs that confer powdery mildew and aphid resistance. Horticulture Research 11, uhad256 (2023).

44. Chovelon, V. et al. Building a cluster of NLR genes conferring resistance to pests and pathogens: the story of the Vat gene cluster in cucurbits. Horticulture research 8, (2021).

45. Chao, A. et al. Rarefaction and extrapolation with Hill numbers: a framework for sampling and estimation in species diversity studies. Ecological Monographs 84, 45–67 (2014).

46. Hsieh, T. C., Ma, K. H. & Chao, A. iNEXT: an R package for rarefaction and extrapolation of species diversity (Hill numbers). Methods in Ecology and Evolution 7, 1451–1456 (2016).

47. Clauw, P., Ellis, T. J., Liu, H.-J. & Sasaki, E. Beyond the Standard GWAS—A Guide for Plant Biologists. Plant and Cell Physiology 66, 431–443 (2025).

48. Oumouloud, A. et al. Characterization of the Fusarium wilt resistance *Fom-2* gene in melon. Molecular Breeding 30, 325–334 (2012).

49. Buerstmayr, M. et al. Fusarium head blight resistance in European winter wheat: insights from genome-wide transcriptome analysis. BMC Genomics 22, 470 (2021).

50. Li, Y. et al. Novel Genomic Regions of Fusarium Wilt Resistance in Bottle Gourd [*Lagenaria siceraria* (Mol.) Standl.] Discovered in Genome-Wide Association Study. Front. Plant Sci. 12, (2021).

51. Kahlon, P. S. & Stam, R. Polymorphisms in plants to restrict losses to pathogens: From gene family expansions to complex network evolution. Current Opinion in Plant Biology 62, 102040 (2021).

52. Mizuno, H. et al. Evolutionary dynamics and impacts of chromosome regions carrying R-gene clusters in rice. Sci Rep 10, 872 (2020).

53. Roman, B., Gomez, P., Pico, B. & Die, J. V. The NBS-LRR Gene Class is a Small Family in *Cucurbita pepo*. Preprint at 10.20944/preprints202001.0048.v1 (2020).

54. Ma, L. et al. Cucurbitaceae genome evolution, gene function, and molecular breeding. Horticulture Research 9, uhab057 (2022).

55. Yang, L. et al. A 1,681-locus consensus genetic map of cultivated cucumber including 67 NB-LRR resistance gene homolog and ten gene loci. BMC Plant Biol 13, 53 (2013).

56. Andolfo, G., Sánchez, C. S., Cañizares, J., Pico, M. B. & Ercolano, M. R. Large-scale gene gains and losses molded the NLR defense arsenal during the Cucurbita evolution. Planta 254, 82 (2021).

57. Song, X. et al. Genome-Wide Analysis of the NBS-LRR Gene Family and SSR Molecular Markers Development in Solanaceae. Horticulturae 10, 1293 (2024).

58. Zhang, Y., Edwards, D. & Batley, J. Comparison and evolutionary analysis of Brassica nucleotide binding site leucine rich repeat (NLR) genes and importance for disease resistance breeding. The Plant Genome 14, e20060 (2021).

59. Saile, S. C. & Kasmi, F. E. Small family, big impact: RNL helper NLRs and their importance in plant innate immunity. PLOS Pathogens 19, e1011315 (2023).

60. Lin, X., Zhang, Y., Kuang, H. & Chen, J. Frequent loss of lineages and deficient duplications accounted for low copy number of disease resistance genes in Cucurbitaceae. BMC Genomics 14, 335 (2013).

61. Bhullar, N. K., Street, K., Mackay, M., Yahiaoui, N. & Keller, B. Unlocking wheat genetic resources for the molecular identification of previously undescribed functional alleles at the *Pm3* resistance locus. Proceedings of the National Academy of Sciences 106, 9519–9524 (2009).

62. Nei, M. Analysis of Gene Diversity in Subdivided Populations. Proceedings of the National Academy of Sciences 70, 3321–3323 (1973).

63. Wehenkel, C., Bergmann, F. & Gregorius, H.R. Is there a trade-off between species diversity and genetic diversity in forest tree communities? Plant Ecol 185, 151–161 (2006).

64. Bergmann, F., Gregorius, H. R., Kownatzki, D. & Wehenkel, C. Different diversity measures and genetic traits reveal different speciesgenetic diversity relationships: A case study in forest tree communities. Silvae Genetica 62, 25–37 (2013).

65. Arora, S. et al. Resistance gene cloning from a wild crop relative by sequence capture and association genetics. Nature biotechnology 37, 139–143 (2019).

66. Ariga, H. et al. NLR locus-mediated trade-off between abiotic and biotic stress adaptation in Arabidopsis. Nature Plants 3, 1–8 (2017).

67. Salinier, J. et al. The INRAE Centre for Vegetable Germplasm: Geographically and Phenotypically Diverse Collections and Their Use in Genetics and Plant Breeding. Plants 11, 347 (2022).

68. Steuernagel, B. et al. The NLR-Annotator Tool Enables Annotation of the Intracellular Immune Receptor Repertoire. Plant Physiol 183, 468–482 (2020).

69. Castanera, R., Ruggieri, V., Pujol, M., Garcia-Mas, J. & Casacuberta, J. M. An Improved Melon Reference Genome With Single-Molecule Sequencing Uncovers a Recent Burst of Transposable Elements With Potential Impact on Genes. Frontiers in Plant Science 10, (2020).

70. Pichot, C. et al. Cantaloupe melon genome reveals 3D chromatin features and structural relationship with the ancestral cucurbitaceae karyotype. iScience 25, 103696 (2022).

71. Yano, R. et al. Comparative genomics of muskmelon reveals a potential role for retrotransposons in the modification of gene expression. Communications Biology 3, (2020).

72. Zhang, H. et al. A High-Quality Melon Genome Assembly Provides Insights into Genetic Basis of Fruit Trait Improvement. iScience 22, 16–27 (2019).

73. Belinchon-Moreno, J. et al. Nuclear and organelle genome assemblies of 5 *Cucumis melo* L. accessions, Ananas, Canton, PI 414723, Vedrantais, and Zhimali, belonging to diverse botanical groups. G3 Genes|Genomes|Genetics 15, jkaf098 (2025).

74. Liu, H., Wu, S., Li, A. & Ruan, J. SMARTdenovo: a *de novo* assembler using long noisy reads. GigaByte 2021, gigabyte15 (2021).

75. Li, H. Minimap2: pairwise alignment for nucleotide sequences. Bioinformatics 34, 3094–3100 (2018).

76. Thorvaldsdóttir, H., Robinson, J. T. & Mesirov, J. P. Integrative Genomics Viewer (IGV): high-performance genomics data visualization and exploration. Briefings in Bioinformatics 14, 178–192 (2013).

77. Guiglielmoni, N., Houtain, A., Derzelle, A., Van Doninck, K. & Flot, J.-F. Overcoming uncollapsed haplotypes in long-read assemblies of non-model organisms. BMC Bioinformatics 22, 303 (2021).

78. Koren, S. et al. Canu: scalable and accurate long-read assembly via adaptive k-mer weighting and repeat separation. Genome Res. 27, 722–736 (2017).

79. Shafin, K. et al. Nanopore sequencing and the Shasta toolkit enable efficient *de novo* assembly of eleven human genomes. Nat Biotechnol 38, 1044–1053 (2020).

80. Hu, J. et al. NextDenovo: an efficient error correction and accurate assembly tool for noisy long reads. Genome Biol 25, 107 (2024).

81. Kolmogorov, M., Yuan, J., Lin, Y. & Pevzner, P. A. Assembly of long, error-prone reads using repeat graphs. Nat Biotechnol 37, 540–546 (2019).

82. Stanojević, D., Lin, D., Sessions, P. F. de & Šikić, M. Telomere-to-telomere phased genome assembly using error-corrected Simplex nanopore reads. Preprint at 10.1101/2024.05.18.594796 (2024).

83. Cheng, H., Concepcion, G. T., Feng, X., Zhang, H. & Li, H. Haplotype-resolved de novo assembly using phased assembly graphs with hifiasm. Nat Methods 18, 170–175 (2021).

84. Danecek, P. et al. Twelve years of SAMtools and BCFtools. GigaScience 10, giab008 (2021).

85. Holst, F. et al. Helixer–de novo Prediction of Primary Eukaryotic Gene Models Combining Deep Learning and a Hidden Markov Model. Preprint at 10.1101/2023.02.06.527280 (2023).

86. Li, P. et al. RGAugury: a pipeline for genome-wide prediction of resistance gene analogs (RGAs) in plants. BMC Genomics 17, 852 (2016).

87. Emms, D. M. & Kelly, S. OrthoFinder: phylogenetic orthology inference for comparative genomics. Genome Biology 20, 238 (2019).

88. Kalinowski, S. T. Counting Alleles with Rarefaction: Private Alleles and Hierarchical Sampling Designs. Conservation Genetics 5, 539–543 (2004).

89. Chao, A., & Chiu, C. H. Species richness: Estimation and comparison: Wiley StatsRef: Statistics Reference Online (John Wiley & Sons, Ltd, 2016).

90. Xia, Y. & Sun, J. Alpha Diversity: Bioinformatic and Statistical Analysis of Microbiome Data (Springer International Publishing, 2023).

91. Bates, D., Mächler, M., Bolker, B. & Walker, S. Fitting Linear Mixed-Effects Models using lme4. Preprint at 10.48550/arXiv.1406.5823 (2014).

92. Perchepied, L., Kroj, T., Tronchet, M., Loudet, O. & Roby, D. Natural Variation in Partial Resistance to Pseudomonas syringae Is Controlled by Two Major QTLs in Arabidopsis thaliana. PLOS ONE 1, e123 (2006).

93. Boissot, N., Thomas, S., Chovelon, V. & Lecoq, H. NBS-LRR-mediated resistance triggered by aphids: viruses do not adapt; aphids adapt via different mechanisms. BMC Plant Biology 16, 25 (2016).

94. Purcell, S. et al. PLINK: A Tool Set for Whole-Genome Association and Population-Based Linkage Analyses. The American Journal of Human Genetics 81, 559–575 (2007).

95. Pritchard, J. K., Stephens, M. & Donnelly, P. Inference of Population Structure Using Multilocus Genotype Data. Genetics 155, 945–959 (2000).

96. Evanno, G., Regnaut, S. & Goudet, J. Detecting the number of clusters of individuals using the software STRUCTURE: a simulation study. Mol Ecol 14, 2611–2620 (2005).

97. Earl, D. A. & vonHoldt, B. M. STRUCTURE HARVESTER: a website and program for visualizing STRUCTURE output and implementing the Evanno method. Conservation Genet Resour 4, 359– 361 (2012).

98. Endelman, J. B. Ridge Regression and Other Kernels for Genomic Selection with R Package rrBLUP. The Plant Genome 4, 250–255 (2011).

99. Hickey, G. et al. Genotyping structural variants in pangenome graphs using the vg toolkit. Genome Biology 21, 35 (2020).

100. Kokot, M., Długosz, M. & Deorowicz, S. KMC 3: counting and manipulating k-mer statistics. Bioinformatics 33, 2759–2761 (2017).

101. Segura, V. et al. An efficient multi-locus mixed-model approach for genome-wide association studies in structured populations. Nat Genet 44, 825–830 (2012).

102. Chen, H. et al. Control for Population Structure and Relatedness for Binary Traits in Genetic Association Studies via Logistic Mixed Models. The American Journal of Human Genetics 98, 653– 666 (2016).

103. Turner, S. D. qqman: an R package for visualizing GWAS results using Q-Q and manhattan plots. 005165 Preprint at 10.1101/005165 (2014).

104. Gao, X. Multiple testing corrections for imputed SNPs. Genetic epidemiology 35, 154 (2011).

105. Mangin, B. et al. Novel measures of linkage disequilibrium that correct the bias due to population structure and relatedness. Heredity 108, 285–291 (2012).

106. Chikhi, R. & Rizk, G. Space-efficient and exact de Bruijn graph representation based on a Bloom filter. Algorithms Mol Biol 8, 22 (2013).

107. Quinlan, A. R. & Hall, I. M. BEDTools: a flexible suite of utilities for comparing genomic features. Bioinformatics 26, 841–842 (2010).

108. Shumate, A. & Salzberg, S. L. Liftoff: accurate mapping of gene annotations. Bioinformatics 37, 1639–1643 (2021).

109. Katoh, K. & Standley, D. M. MAFFT Multiple Sequence Alignment Software Version 7: Improvements in Performance and Usability. Molecular Biology and Evolution 30, 772–780 (2013).

110. Nguyen, L. T., Schmidt, H. A., von Haeseler, A. & Minh, B. Q. IQ-TREE: A Fast and Effective Stochastic Algorithm for Estimating Maximum-Likelihood Phylogenies. Molecular Biology and Evolution 32, 268–274 (2015).

111. Letunic, I. & Bork, P. Interactive Tree of Life (iTOL) v6: recent updates to the phylogenetic tree display and annotation tool. Nucleic Acids Research 52, W78–W82 (2024).

112. Cabanettes, F. & Klopp, C. D-GENIES: dot plot large genomes in an interactive, efficient and simple way. PeerJ 6, e4958 (2018).

113. Belinchon-Moreno, J., Boissot, N. & Faivre-Rampant, P. NLRome assemblies of *Cucumis melo* derived from Nanopore adaptive sampling sequencing. Recherche Data Gouv Dataset 10.57745/ZALVPU (2025).

114. Belinchon-Moreno, J., Boissot, N. & Faivre-Rampant, P. Integrated genotypic and phenotypic dataset for multi-pathogen resistance GWAS in *Cucumis melo*. Recherche Data Gouv Dataset 10.57745/AUO6YY (2025).

