## Extended Data Figures for "Intra-specific NLR allelic diversity and genomic landscape for plant resistance association studies"

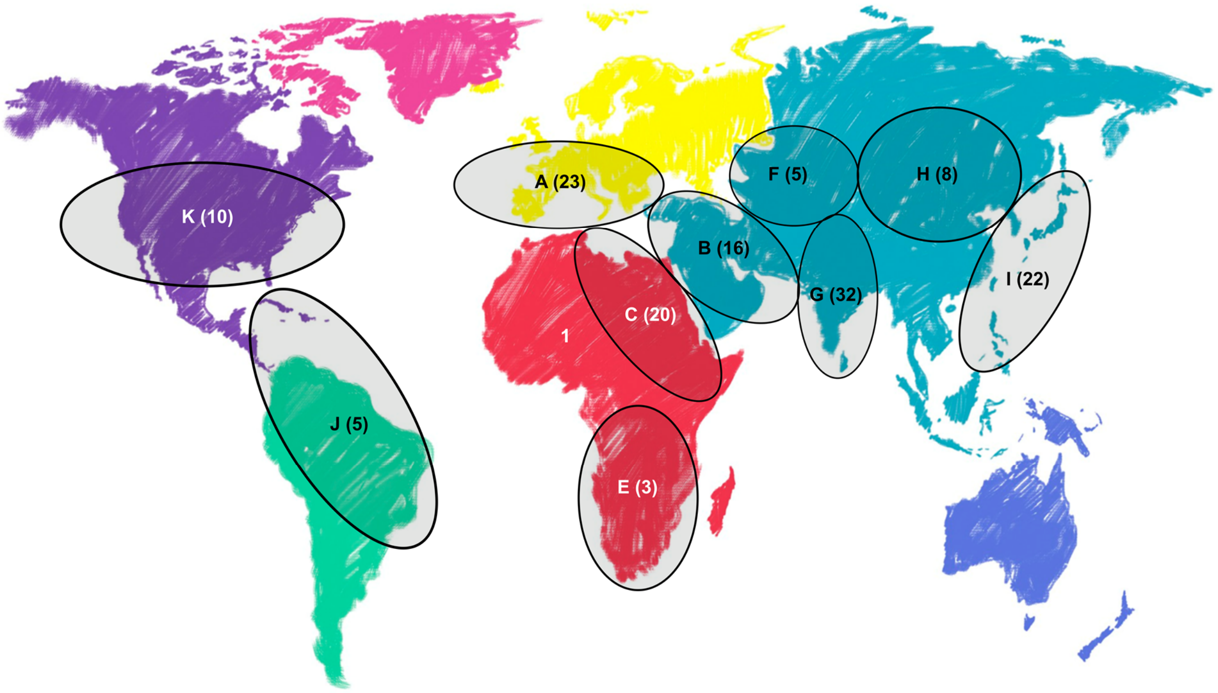


***Extended Data Fig. 1.*** *Geographical origin of the panel of 151 accessions sequenced with NAS. Each geographical origin is represented by a distinct letter, with the number of accessions from each origin indicated in parentheses. Six accessions are not shown, as their geographical origin is unknown.*

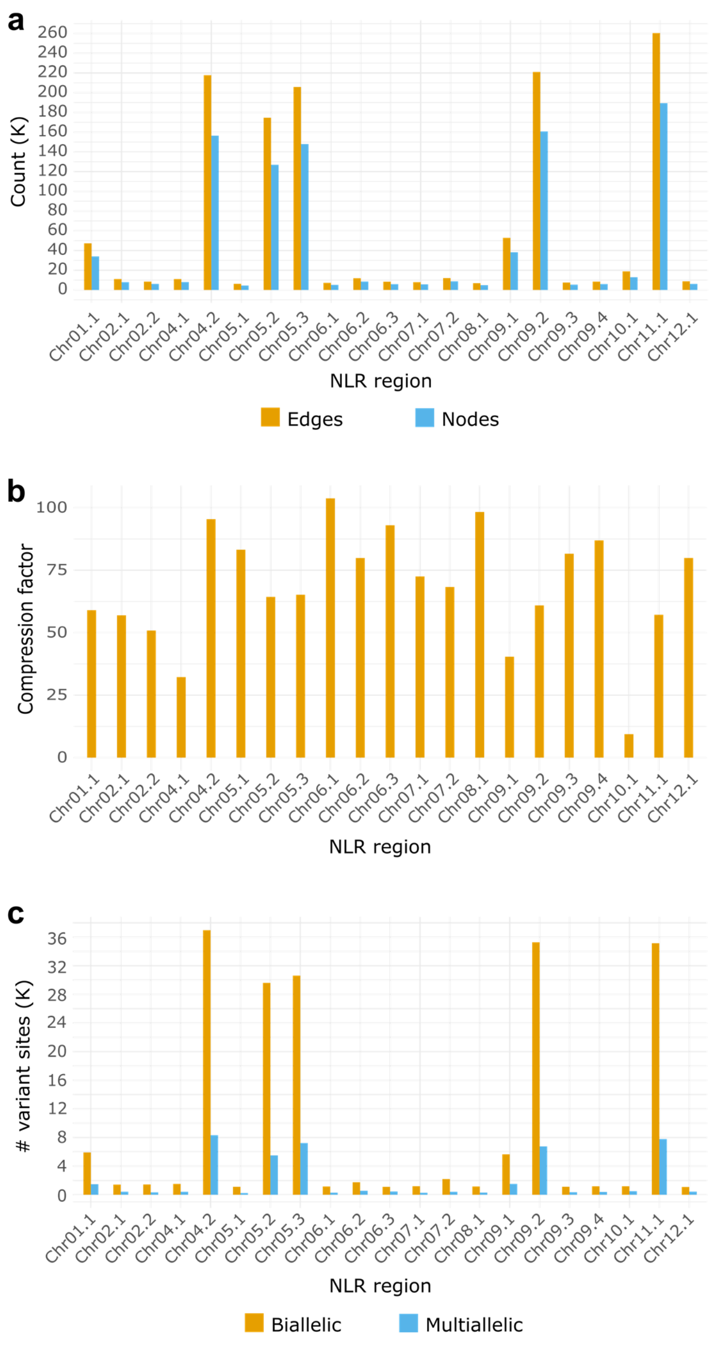


***Extended Data Fig. 2.*** *Metrics of the pan-NLRome variation graph. The pan-NLRome graph was composed of 21 sub-graphs (one for each NLR region) and constructed from the assembled haplotypes of 143 accessions.* ***a,*** *Distribution of nodes and edges counts for each pan-NLRome sub-graphs.* ***b,*** *Compression factor of each pan-NLRome sub-graph, calculated as the total size of all genomic sequences included in the sub-graph divided by the final sub-graph size.* ***c,*** *Distribution of the number of biallelic and multiallelic variant sites across the 21 pan-NLRome sub-graphs. Variants were identified using the ANSO-77 accession as the reference path through each graph.*

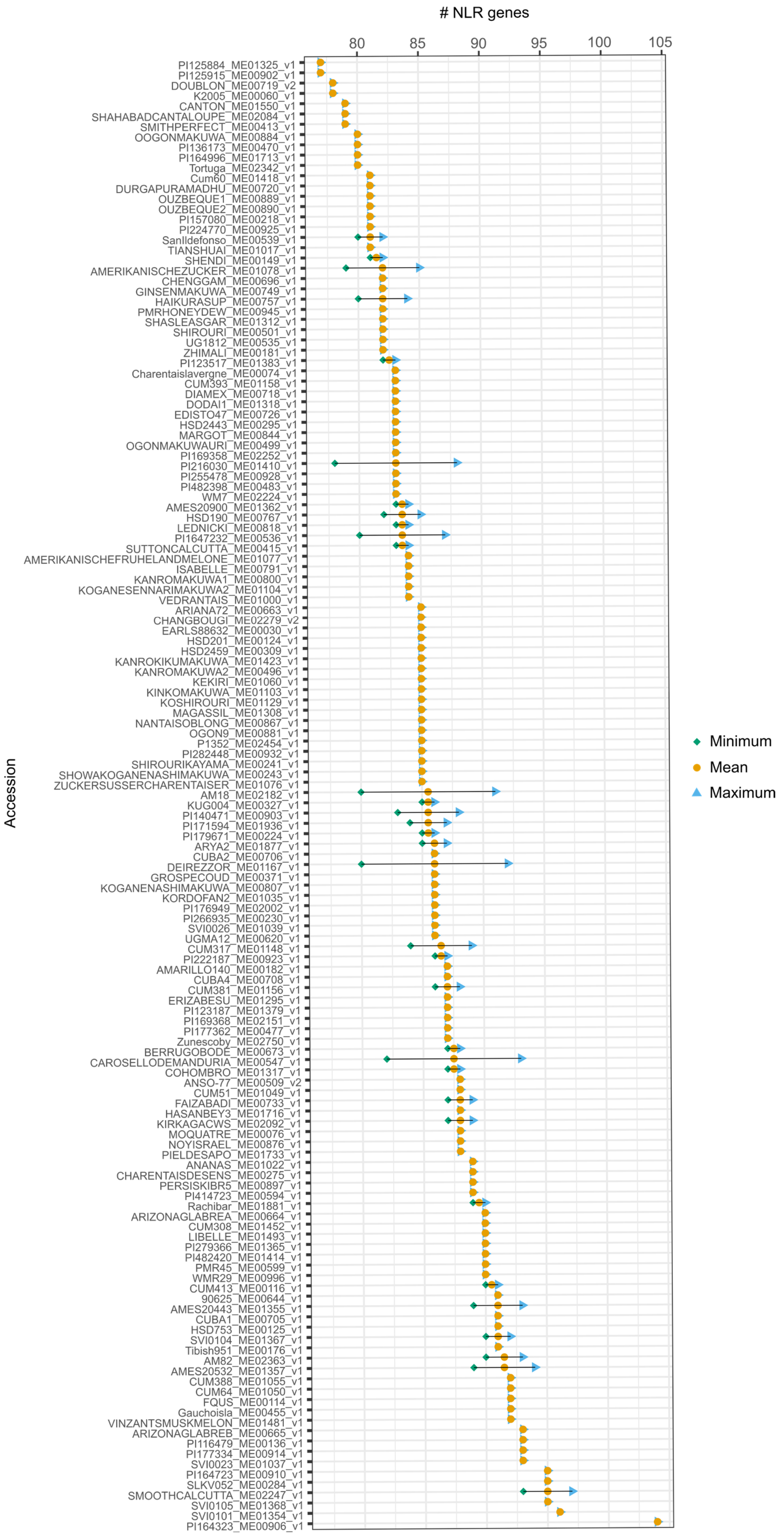

### NLR genes
80
85
90
95
100
105
Accession
PI125884_ME01325_v1
PI125915_ME00902_v1
DOUBLON_ME00719_v2
K2005_ME00060_v1
CANTON_ME01550_v1
SHAHABADCANTALOUPE_ME02084_v1
SMITHPERFECT_ME00413_v1
OOGONMAKUWA_ME00884_v1
PI136173_ME00470_v1
PI164996_ME01713_v1
Tortuga_ME02342_v1
Cum60_ME01418_v1
DURGAPURAMADHU_ME00720_v1
OUZBEQUE1_ME00889_v1
OUZBEQUE2_ME00890_v1
PI157080_ME00218_v1
PI224770_ME00925_v1
SanIldefonso_ME00539_v1
TIANSHUAI_ME01017_v1
SHENDI_ME00149_v1
AMERIKANISCHEZUCKER_ME01078_v1
CHENGGAM_ME00696_v1
GINSENMAKUWA_ME00749_v1
HAIKURASUP_ME00757_v1
PMRHONEYDEW_ME00945_v1
SHASLEASGAR_ME01312_v1
SHIROURI_ME00501_v1
UG1812_ME00535_v1
ZHIMALI_ME00181_v1
PI123517_ME01383_v1
Charentaislavergne_ME00074_v1
CUM393_ME01158_v1
DIAMEX_ME00718_v1
DODAI1_ME01318_v1
EDISTO47_ME00726_v1
HSD2443_ME00295_v1
MARGOT_ME00844_v1
OGONMAKUWAURI_ME00499_v1
PI169358_ME02252_v1
PI216030_ME01410_v1
PI255478_ME00928_v1
PI482398_ME00483_v1
WM7_ME02224_v1
AMES20900_ME01362_v1
HSD190_ME00767_v1
LEDNICKI_ME00818_v1
PI1647232_ME00536_v1
SUTTONCALCUTTA_ME00415_v1
AMERIKANISCHEFRUHELANDMELONE_ME01077_v1
ISABELLE_ME00791_v1
KANROMAKUWA1_ME00800_v1
KOGANESENNARIMAKUWA2_ME01104_v1
VEDRANTAIS_ME01000_v1
ARIANA72_ME00663_v1
CHANGBOUGI_ME02279_v2
EARLS88632_ME00030_v1
HSD201_ME00124_v1
HSD2459_ME00309_v1
KANROKIKUMAKUWA_ME01423_v1
KANROMAKUWA2_ME00496_v1
KEKIRI_ME01060_v1
KINKOMAKUWA_ME01103_v1
KOSHIROURI_ME01129_v1
MAGASSIL_ME01308_v1
NANTAISOBLONG_ME00867_v1
OGON9_ME00881_v1
P1352_ME02454_v1
PI282448_ME00932_v1
SHIROURIKAYAMA_ME00241_v1
SHOWAKOGANENASHIMAKUWA_ME00243_v1
ZUCKERSUSSERCHARENTAISER_ME01076_v1
AM18_ME02182_v1
KUG004_ME00327_v1
PI140471_ME00903_v1
PI171594_ME01936_v1
PI179671_ME00224_v1
ARYA2_ME01877_v1
CUBA2_ME00706_v1
DEIREZZOR_ME01167_v1
GROSPECOUD_ME00371_v1
KOGANENASHIMAKUWA_ME00807_v1
KORDOFAN2_ME01035_v1
PI176949_ME02002_v1
PI266935_ME00230_v1
SVI0026_ME01039_v1
UGMA12_ME00620_v1
CUM317_ME01148_v1
PI222187_ME00923_v1
AMARILLO140_ME00182_v1
CUBA4_ME00708_v1
CUM381_ME01156_v1
ERIZABESU_ME01295_v1
PI123187_ME01379_v1
PI169368_ME02151_v1
PI177362_ME00477_v1
Zunescoby_ME02750_v1
BERRUGOBODE_ME00673_v1
CAROSELLODEMANDURIA_ME00547_v1
COHOMBRO_ME01317_v1
ANSO-77_ME00509_v2
CUM51_ME01049_v1
FAIZABADI_ME00733_v1
HASANBEY3_ME01716_v1
KIRKAGACWS_ME02092_v1
MOQUATRE_ME00076_v1
NOYISRAEL_ME00876_v1
PIELDESAPO_ME01733_v1
ANANAS_ME01022_v1
CHARENTAISDESENS_ME00275_v1
PERSISKIBR5_ME00897_v1
PI414723_ME00594_v1
Rachibar_ME01881_v1
ARIZONAGLABREA_ME00664_v1
CUM308_ME01452_v1
LIBELLE_ME01493_v1
PI279366_ME01365_v1
PI482420_ME01414_v1
PMR45_ME00599_v1
WMR29_ME00996_v1
CUM413_ME00116_v1
90625_ME00644_v1
AMES20443_ME01355_v1
CUBA1_ME00705_v1
HSD753_ME00125_v1
SVI0104_ME01367_v1
Tibish951_ME00176_v1
AM82_ME02363_v1
AMES20532_ME01357_v1
CUM388_ME01055_v1
CUM64_ME01050_v1
FQUS_ME00114_v1
Gauchoisla_ME00455_v1
VINZANTSMUSKMELON_ME01481_v1
ARIZONAGLABREB_ME00665_v1
PI116479_ME00136_v1
PI177334_ME00914_v1
SVI0023_ME01037_v1
PI164723_ME00910_v1
SLKV052_ME00284_v1
SMOOTHCALCUTTA_ME02247_v1
SVI0105_ME01368_v1
SVI0101_ME01354_v1
PI164323_ME00906_v1
Minimum
Mean
Maximum

***Extended Data Fig. 3.*** *Minimum, maximum, and average NLR content per sequenced and assembled accession. For accessions in which certain NLR regions were assembled into two pseudo-haplotypes, both the minimum and maximum possible NLR counts were estimated based on the structure of these pseudo-haplotypes.*

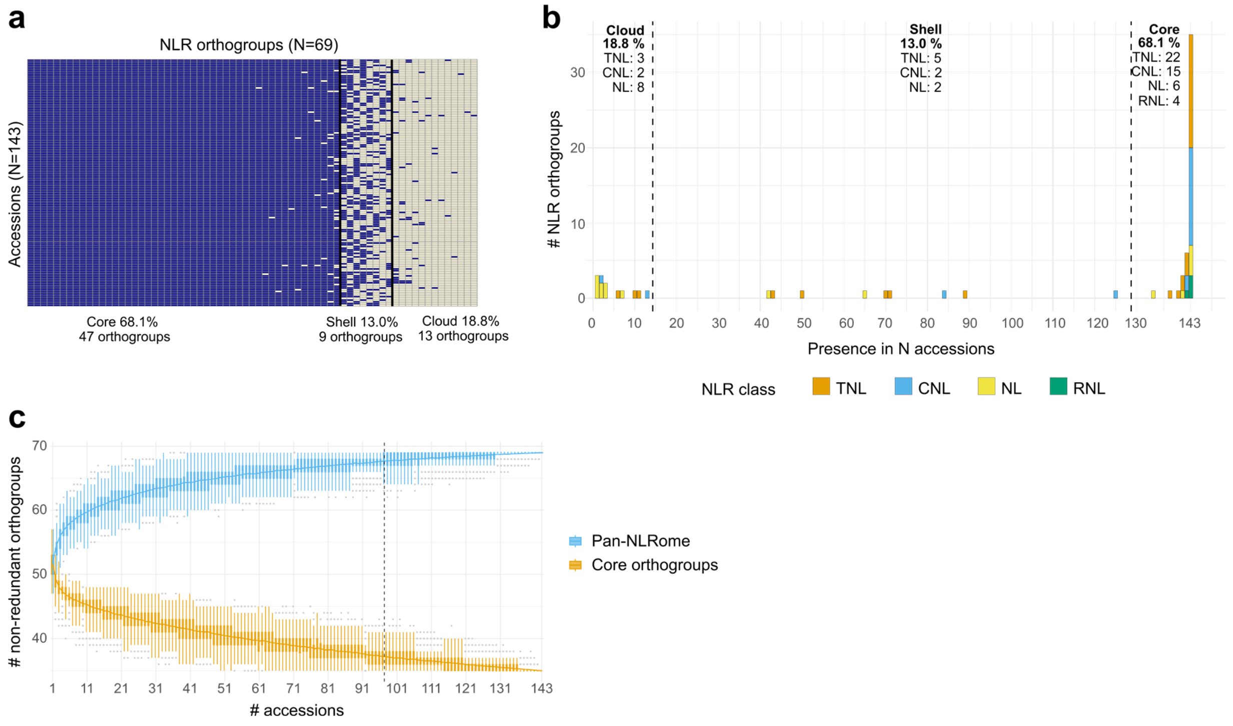


***Extended Data Fig. 4.*** *Orthogroup-based saturation analysis of the pan-NLRome. Orthogroups were obtained using the orthofinder software with default parameters, providing as input the annotated NLR alleles from the 143 sequenced and assembled accessions. Four orthogroups were removed as they corresponded to single, truncated alleles.* ***a****, Heatmap representing the variation patterns of the inferred NLR orthogroups across the 143 accessions. Blue squares represent NLR orthogroups present in the corresponding accession, while grey squares represent those absent. Two black vertical line divides NLR orthogroups belonging to the core (≥90% of accessions), shell (10%≤N<90%) and cloud (<10% of accessions) pan-genome.* ***b,*** *Distribution of the number of NLR orthogroups shared by N accessions among different NLR classes in the cloud, shell and core pan-genome. The NLR classes of NLR orthogroups were defined as the most represented class among NLR alleles composing each NLR orthogroup. The percentage of NLR orthogroups belonging to the cloud, shell and core pan-genome is shown at the top, with the detailed number of NLR orthogroups per class displayed below it.* ***c,*** *Saturation growth curve of the variation in the number of NLR orthogroups within the pan-NLRome and core NLRome. Boxplots were fitted at each given number of accessions, with a colored line connecting medians. Grey dots mark outliers beyond 1.5 x Interquartile Ratio, and whiskers cover all non-outlier values. A black, dotted, vertical line mark the minimum number of accessions needed to obtain 95% of the observed NLR orthogroups.*

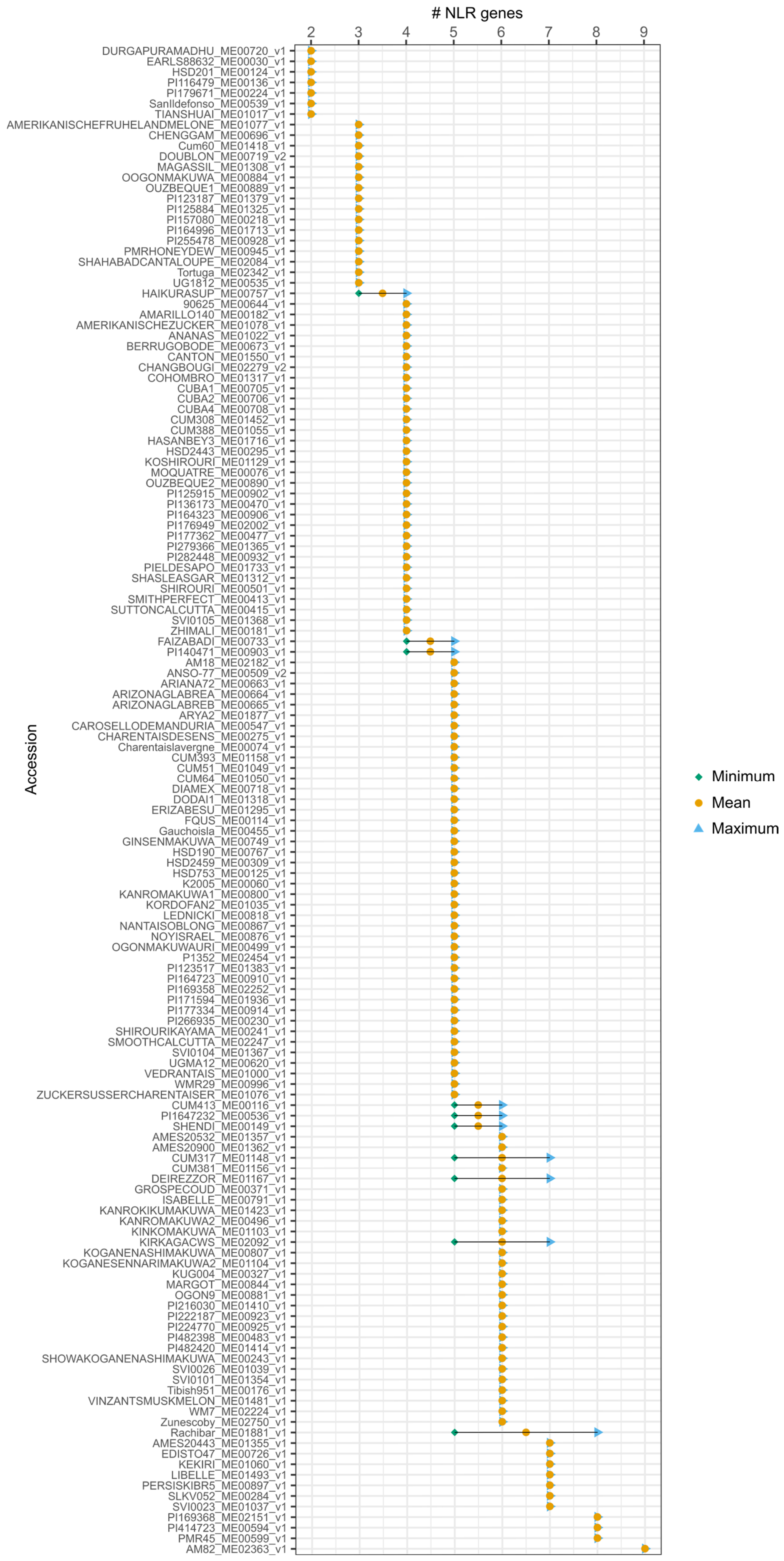

### NLR genes
2
3
4
5
6
7
8
9
Accession
DURGAPURAMADHU_ME00720_v1
EARLS88632_ME00030_v1
HSD201_ME00124_v1
PI116479_ME00136_v1
PI179671_ME00224_v1
SanIldefonso_ME00539_v1
TIANSHUAI_ME01017_v1
AMERIKANISCHEFRUHELANDMELONE_ME01077_v1
CHENGGAM_ME00696_v1
Cum60_ME01418_v1
DOUBLON_ME00719_v2
MAGASSIL_ME01308_v1
OOGONMAKUWA_ME00884_v1
OUZBEQUE1_ME00889_v1
PI123187_ME01379_v1
PI125884_ME01325_v1
PI157080_ME00218_v1
PI164996_ME01713_v1
PI255478_ME00928_v1
PMRHONEYDEW_ME00945_v1
SHAHABADCANTALOUPE_ME02084_v1
Tortuga_ME02342_v1
UG1812_ME00535_v1
HAIKURASUP_ME00757_v1
90625_ME00644_v1
AMARILLO140_ME00182_v1
AMERIKANISCHEZUCKER_ME01078_v1
ANANAS_ME01022_v1
BERRUGOBODE_ME00673_v1
CANTON_ME01550_v1
CHANGBOUGI_ME02279_v2
COHOMBRO_ME01317_v1
CUBA1_ME00705_v1
CUBA2_ME00706_v1
CUBA4_ME00708_v1
CUM308_ME01452_v1
CUM388_ME01055_v1
HASANBEY3_ME01716_v1
HSD2443_ME00295_v1
KOSHIROURI_ME01129_v1
MOQUATRE_ME00076_v1
OUZBEQUE2_ME00890_v1
PI125915_ME00902_v1
PI136173_ME00470_v1
PI164323_ME00906_v1
PI176949_ME02002_v1
PI177362_ME00477_v1
PI279366_ME01365_v1
PI282448_ME00932_v1
PIELDESAPO_ME01733_v1
SHASLEASGAR_ME01312_v1
SHIROURI_ME00501_v1
SMITHPERFECT_ME00413_v1
SUTTONCALCUTTA_ME00415_v1
SVI0105_ME01368_v1
ZHIMALI_ME00181_v1
FAIZABADI_ME00733_v1
PI140471_ME00903_v1
AM18_ME02182_v1
ANSO-77_ME00509_v2
ARIANA72_ME00663_v1
ARIZONAGLABREA_ME00664_v1
ARIZONAGLABREB_ME00665_v1
ARYA2_ME01877_v1
CAROSELLODEMANDURIA_ME00547_v1
CHARENTAISDESENS_ME00275_v1
Charentaislavergne_ME00074_v1
CUM393_ME01158_v1
CUM51_ME01049_v1
CUM64_ME01050_v1
DIAMEX_ME00718_v1
DODAI1_ME01318_v1
ERIZABESU_ME01295_v1
FQUS_ME00114_v1
Gauchoisla_ME00455_v1
GINSENMAKUWA_ME00749_v1
HSD190_ME00767_v1
HSD2459_ME00309_v1
HSD753_ME00125_v1
K2005_ME00060_v1
KANROMAKUWA1_ME00800_v1
KORDOFAN2_ME01035_v1
LEDNICKI_ME00818_v1
NANTAISOBLONG_ME00867_v1
NOYISRAEL_ME00876_v1
OGONMAKUWAURI_ME00499_v1
P1352_ME02454_v1
PI123517_ME01383_v1
PI164723_ME00910_v1
PI169358_ME02252_v1
PI171594_ME01936_v1
PI177334_ME00914_v1
PI266935_ME00230_v1
SHIROURIKAYAMA_ME00241_v1
SMOOTHCALCUTTA_ME02247_v1
SVI0104_ME01367_v1
UGMA12_ME00620_v1
VEDRANTAIS_ME01000_v1
WMR29_ME00996_v1
ZUCKERSUSSERCHARENTAISER_ME01076_v1
CUM413_ME00116_v1
PI1647232_ME00536_v1
SHENDI_ME00149_v1
AMES20532_ME01357_v1
AMES20900_ME01362_v1
CUM317_ME01148_v1
CUM381_ME01156_v1
DEIREZZOR_ME01167_v1
GROSPECOUD_ME00371_v1
ISABELLE_ME00791_v1
KANROKIKUMAKUWA_ME01423_v1
KANROMAKUWA2_ME00496_v1
KINKOMAKUWA_ME01103_v1
KIRKAGACWS_ME02092_v1
KOGANENASHIMAKUWA_ME00807_v1
KOGANESENNARIMAKUWA2_ME01104_v1
KUG004_ME00327_v1
MARGOT_ME00844_v1
OGON9_ME00881_v1
PI216030_ME01410_v1
PI222187_ME00923_v1
PI224770_ME00925_v1
PI482398_ME00483_v1
PI482420_ME01414_v1
SHOWAKOGANENASHIMAKUWA_ME00243_v1
SVI0026_ME01039_v1
SVI0101_ME01354_v1
Tibish951_ME00176_v1
VINZANTSMUSKMELON_ME01481_v1
WM7_ME02224_v1
Zunescoby_ME02750_v1
Rachibar_ME01881_v1
AMES20443_ME01355_v1
EDISTO47_ME00726_v1
KEKIRI_ME01060_v1
LIBELLE_ME01493_v1
PERSISKIBR5_ME00897_v1
SLKV052_ME00284_v1
SVI0023_ME01037_v1
PI169368_ME02151_v1
PI414723_ME00594_v1
PMR45_ME00599_v1
AM82_ME02363_v1
Minimum
Mean
Maximum

***Extended Data Fig. 5***. *Minimum, maximum, and average NLR content per accession within the Vat region. For accessions in which the Vat region was assembled into two pseudo-haplotypes, both the minimum and maximum possible NLR counts were estimated based on the structure of these pseudo-haplotypes.*

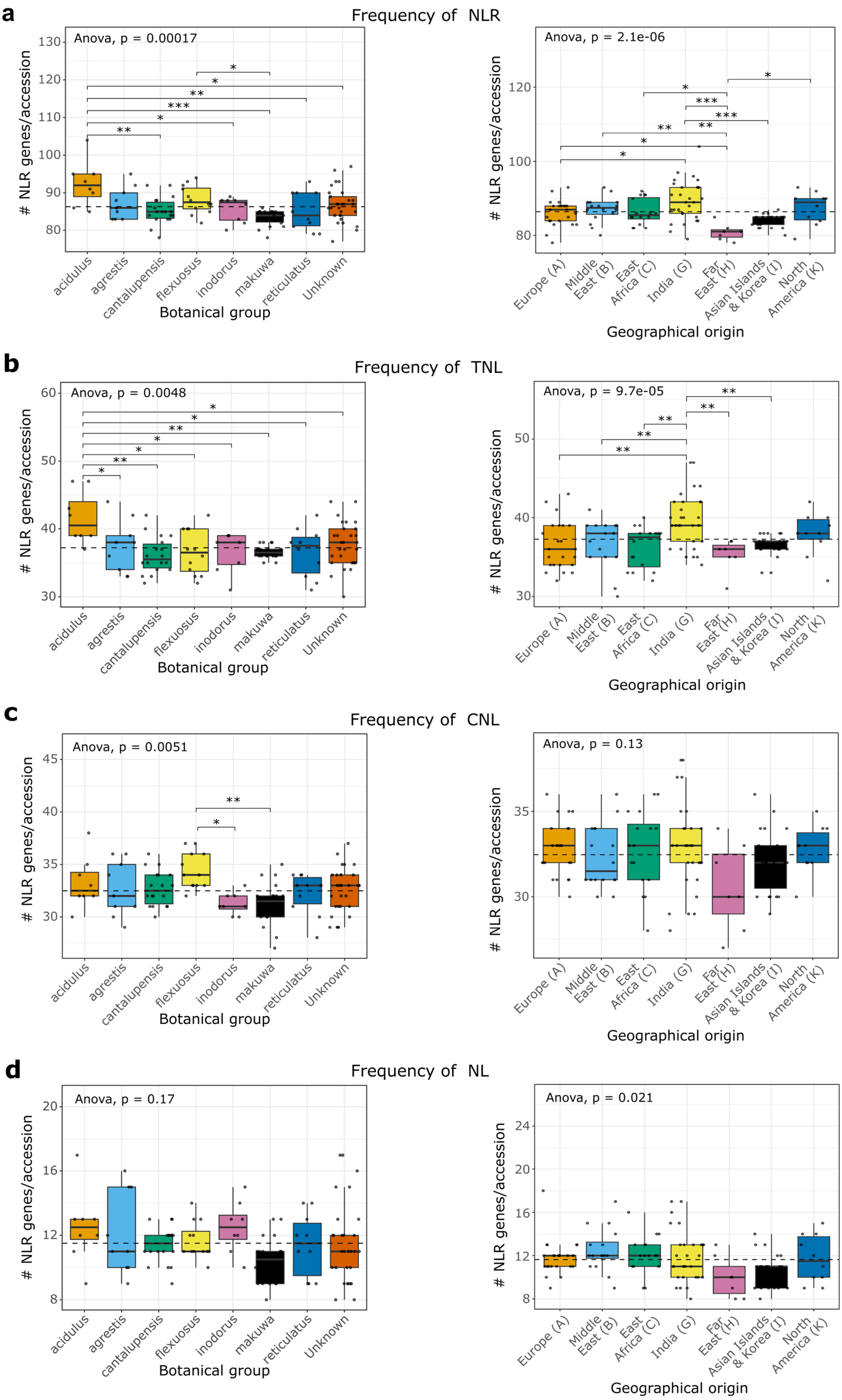

a
Frequency of NLR
Anova, p = 0.00017
Anova, p = 2.1e-06
b
Frequency of TNL
Anova, p = 0.0048
Anova, p = 9.7e-05
c
Frequency of CNL
Anova, p = 0.0051
Anova, p = 0.13
d
Frequency of NL
Anova, p = 0.17
Anova, p = 0.021
### NLR genes/accession
acidulus
agrestis
cantalupensis
flexuosus
inodorus
makuwa
reticulatus
Unknown
Botanical group
Europe (A)
Middle East (B)
East Africa (C)
India (G)
Far East (H)
Asian Islands & Korea (I)
North America (K)
Geographical origin

***Extended Data Fig. 6.*** *NLR content distribution across botanical groups and geographical origins by NLR class.* ***a****, Total number of NLRs.* ***b****, TNLs.* ***c****, CNLs.* ***d****, NLs. For accessions with more than one pseudo-haplotype assembled in any NLR region, the path with the highest NLR count was selected. Only botanical or geographical groups with at least seven accessions were included in the analysis. A one-way analysis of variance (ANOVA) was performed to assess whether the mean number of NLRs per accession differed significantly among botanical groups or geographical origins. The ANOVA p-values are displayed in the top left corner of each plot. For comparisons showing significant ANOVA results, a post-hoc Tukey's HSD test was performed at a 95% confidence level to identify specific group-wise differences. Significance levels are indicated as follows: (*) p-value $< 0.05$; (**) p-value $< 0.05$; (***) p-value $< 0.01$. Solid black horizontal lines inside box-plots represent group medians; small black dots represent individual accession values. Dashed horizontal lines indicate the overall mean NLR content across all groups.*

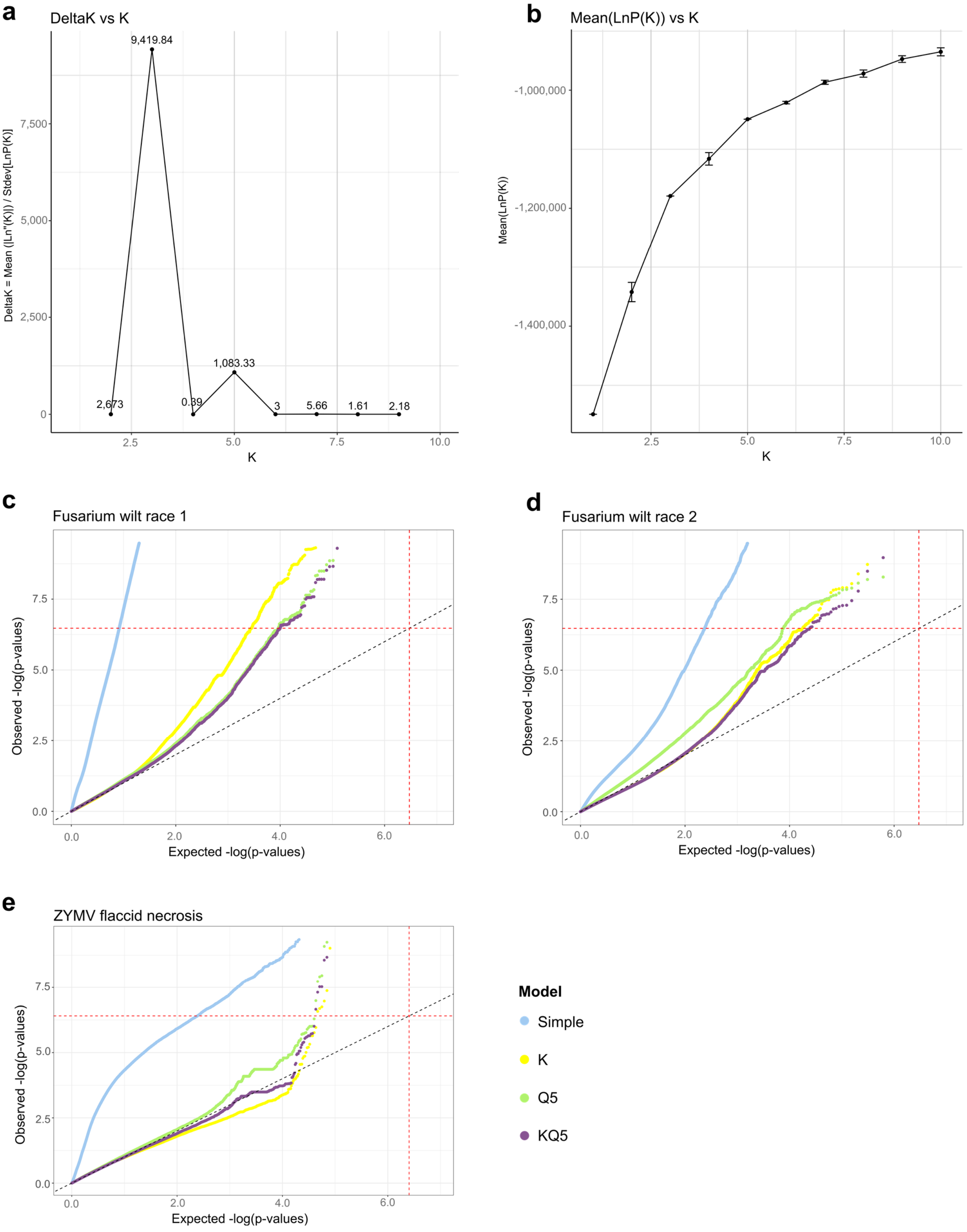


***Extended Data Fig. 7****. Evanno plots derived from STRUCTURE analysis for population structure inference and quantile-quantile (Q-Q) plots from SNP-based GWAS for model selection.* ***a****, Evanno's Delta K statistics for ten tested values of K (number of genetic clusters), each with ten replicates. Correlation between K and Delta K suggested that the population is most likely structured into three or five genetic groups.* ***b****, Estimation of population structure*

*based on the mean values of LnP(K) (log probability of data) plus/minus standard deviation across ten replicates for each K value. Similar to (a), this test supported three or five as the most likely number of genetically distinct populations. c, Q-Q plot of GWAS for Fusarium wilt race 1. d, Q-Q plot of GWAS for Fusarium wilt race 2. e, Q-Q plot of GWAS for ZYMV flaccid necrosis. All Q-Q plots are based on the random regression model. Colored dots indicate –log10(p-values) of SNPs tested under different models. The diagonal dashed line represents the expected distribution of –log10(p-values) under the null hypothesis of no association. Red horizontal and vertical dashed lines indicate the Bonferroni-corrected significance threshold, calculated as α=0.05 divided by the estimated number of independent SNPs.*

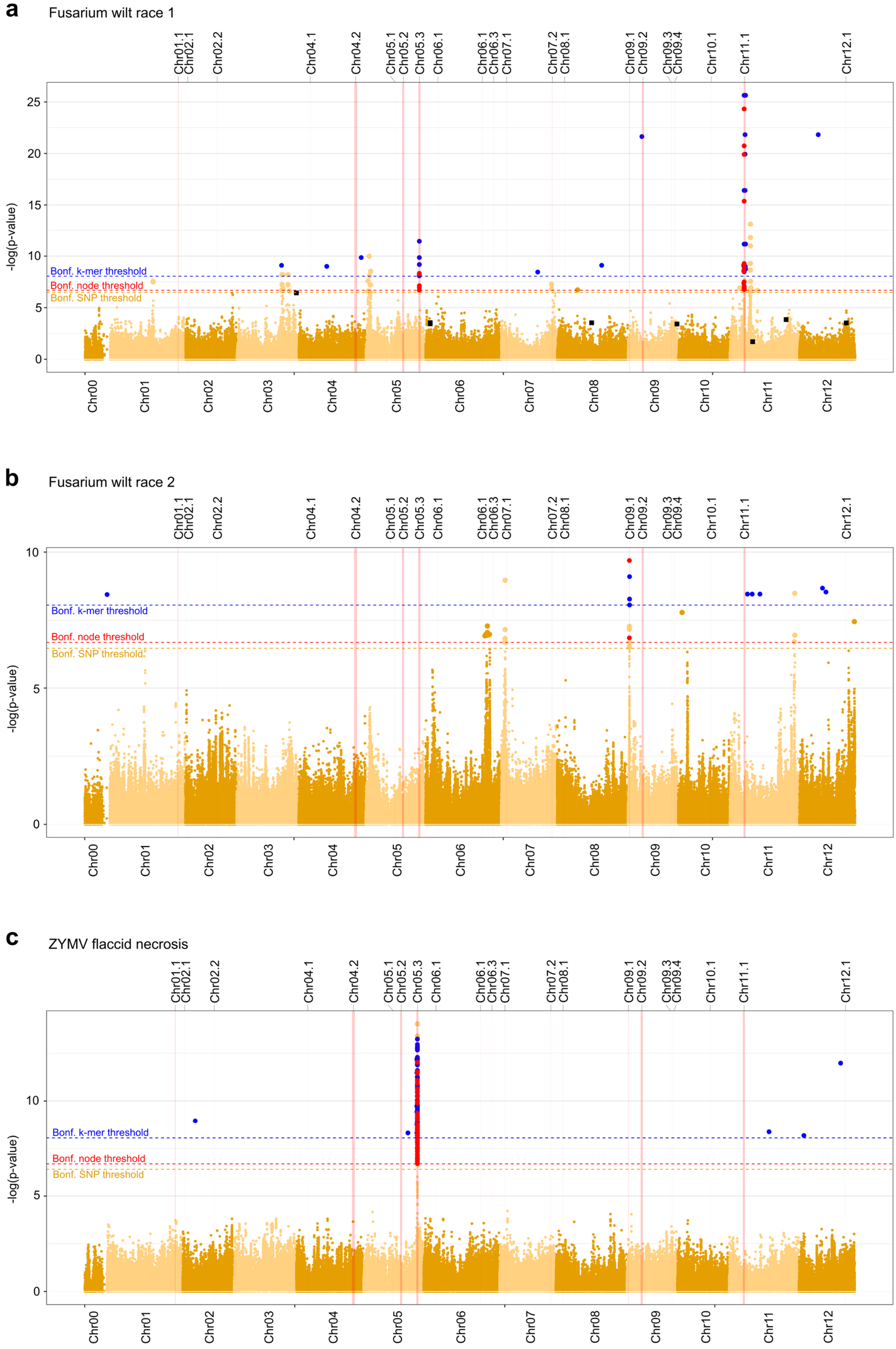
a
Fusarium wilt race 1
b
Fusarium wilt race 2
c
ZYMV flaccid necrosis
Chr01.1
Chr02.1
Chr02.2
Chr04.1
Chr04.2
Chr05.1
Chr05.2
Chr05.3
Chr06.1
Chr06.3
Chr07.1
Chr07.2
Chr08.1
Chr09.1
Chr09.2
Chr09.3
Chr09.4
Chr10.1
Chr11.1
Chr12.1
Bonf. k-mer threshold
Bonf. node threshold
Bonf. SNP threshold
-log(p-value)
Chr00
Chr01
Chr02
Chr03
Chr04
Chr05
Chr06
Chr07
Chr08
Chr09
Chr10
Chr11
Chr12

***Extended Data Fig. 8***. *SNP-, graph- and k-mer based GWAS results presented as Manhattan plots.* ***a****, Fusarium wilt race 1.* ***b****, Fusarium wilt race 2.* ***c****, ZYMV flaccid necrosis. All genomic positions are mapped to the ANSO-77 genome assembly. Orange dots represent whole-genome SNPs; blue dots represent significant k-mers; red dots indicate significant graph nodes. Black squares highlight significant SNPs retained in further steps of the Multi-Locus Mixed Model (MLMM). Bonferroni-corrected significance thresholds (α = 0.05, adjusted for the estimated number of independent variants) are indicated by horizontal lines: orange for SNPs, blue for k-mers, and red for graph-based nodes.*

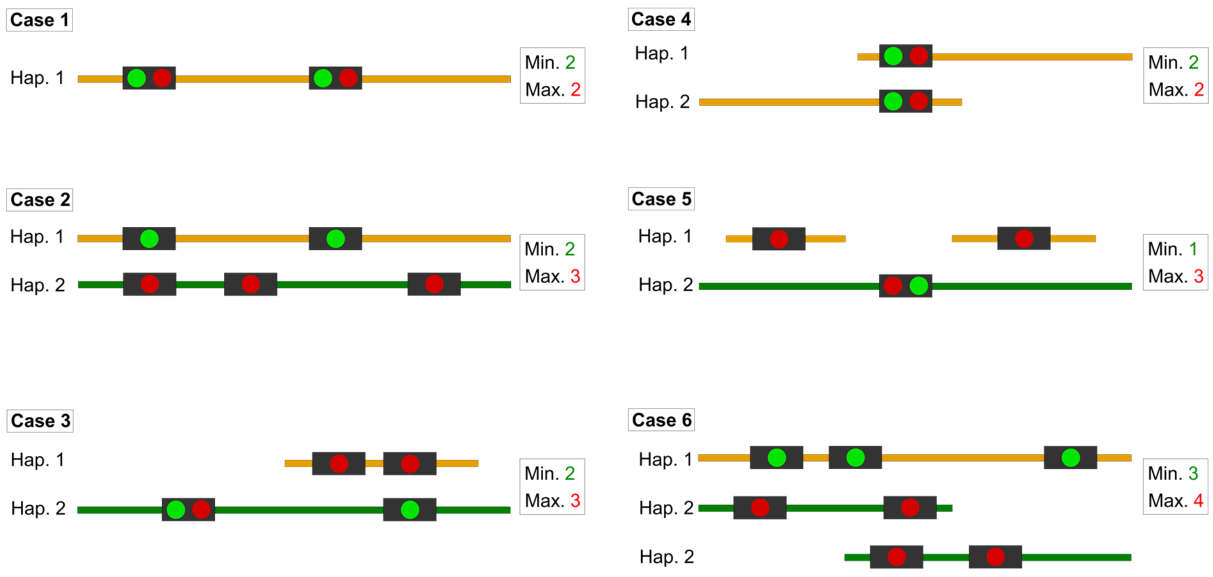


***Extended Data Fig. 9.*** *Cases for the identification of minimum and maximum possible NLR counts per NLR region. Different assembled pseudo-haplotypes are represented by orange and green horizontal bars. NLR genes are shown as black squares. The paths followed to get the minimum (green) and maximum (red) possible NLR count for each region are indicated by green and red circles placed within the corresponding NLR genes. NLR regions with a single haplotype had logically the same minimum and maximum NLR counts (Case 1). In NLR regions with two end-to-end pseudo-haplotypes, we counted the NLRs in each haplotype to determine the maximum and minimum values (Case 2). If one haplotype was not covering the whole NLR region, we included the NLR count from the other haplotype for the non-overlapping region (Cases 3 and 5). When the end of a contig overlapped with the start of another, and NLRs were annotated in the overlapping segment, we counted them in both contigs, as this overlapping could have resulted from the existence of a gene duplication (Cases 4 and 6).*

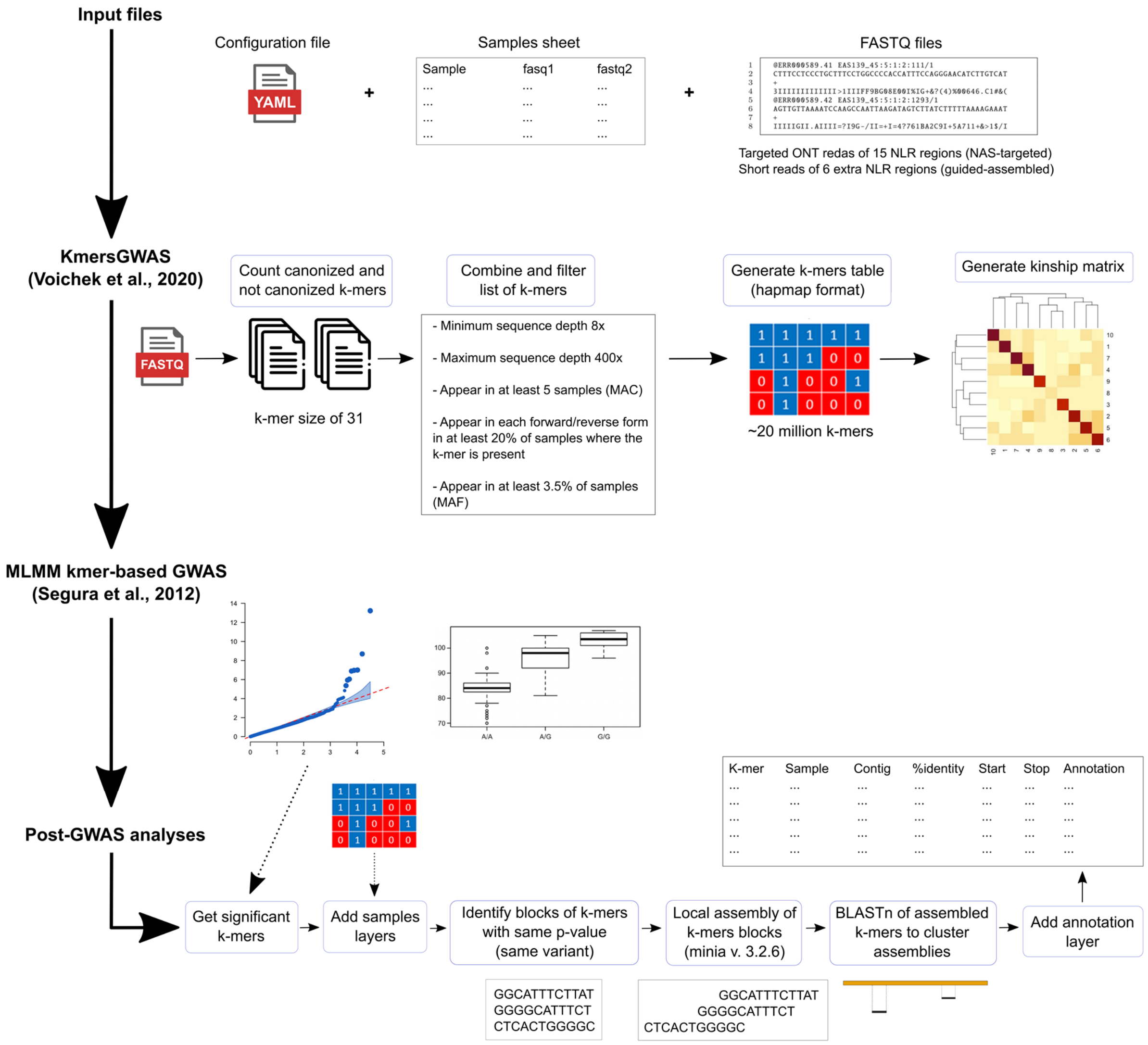


***Extended Data Fig. 10.*** *Overview of the k-mers-based GWAS approach. The method consists on three phases: the extraction of k-mers, k-mer matrix generation and k-mer kinship matrix production (first step), the k-mers-based GWAS (second step), and the post-GWAS analyses (third step). Configuration settings and input files are displayed at the top. The final output file format is displayed in the bottom right section.*
